# Structural Impact of Alzheimer’s Disease Mutations on Transmembrane TREM2-DAP12 Interactions: An Atomistic Perspective

**DOI:** 10.1101/2025.06.26.661739

**Authors:** Zhiwen Zhong, Martin B. Ulmschneider, Christian D. Lorenz

## Abstract

Triggering receptor expressed on myeloid cells 2 (TREM2) is an immunomodulatory receptor that plays a critical role in microglial activation through its association with the adaptor protein DNAX-activation protein 12 (DAP12). Genetic studies have identified rare TREM2 variants as key risk factors for Alzheimer’s disease (AD), with mutations affecting both the extracellular and transmembrane domains. While the TREM2-DAP12 complex is essential for microglial activation and is implicated in Alzheimer’s disease pathology, the structural and mechanistic effects of transmembrane domain mutations, particularly within different TREM2 isoforms, remain unclear. In this study, we employ multi-scale molecular dynamics simulations to investigate the impact of six TREM2 mutations (four in isoform 1, two in isoform 2) and one wild-type control (isoform 2) on complex stability. By integrating coarse-grain and all-atom simulations with unsupervised machine learning techniques, we reveal distinct conformational states and residue-level interactions that govern complex formation. Our findings demonstrate how mutations such as W191X in TREM2 isoform 2 disrupt critical interactions, leading to destabilisation of the complex and potential impairments in neuroimmune signalling. These results provide new insights into the structural mechanisms linking TREM2 mutations to AD and offer a foundation for developing targeted therapeutic strategies.

## INTRODUCTION

Alzheimer’s disease (AD) is a progressive neurodegenerative condition characterised mainly by *β*-amyloid (A*β*) plaque deposition in the brain (Jonsson et al. 2013b; Huang and Mucke 2012). Microglia, the brain’s resident immune cells, are frequently found surrounding A*β* plaques in both AD patients (D’Andrea et al. 2004) and transgenic mouse models (Dickson 1999), where they contribute to the clearance of A*β* and modulation of neuroinflammation. Recent genome-wide association studies (GWAS) have identified rare variants in TREM2 (Triggering Receptor Expressed on Myeloid cells 2) as significant genetic risk factors for late-onset AD, with the R47H mutation in the extracellular domain conferring a particularly strong effect (Guerreiro et al. 2013b; Jonsson et al. 2013a). TREM2 functions as an immunomodulatory receptor on microglial surfaces and initiates intracellular signalling via its transmembrane adaptor DAP12 (DNAX-activating protein of 12 kDa) (Colonna 2023; Ulland and Colonna 2018).

TREM2 interacts with a diverse range of ligands such as phospholipids, sulfatides, bacterial lipopolysaccharide (LPS) and DNA (Wang et al. 2015; Cannon et al. 2012; Daws et al. 2003; Kawabori et al. 2015), and the nature of these interactions can differentially modulate its signalling strength and outcome (Kober and Brett 2017). Ligand engagement at the TREM2 extracellular domain initiates the signalling cascade through association with adaptor proteins DAP10/12 (Ulland et al. 2017). Subsequently, the TREM2 ectodomain is shed by ADAM10/17 metalloproteases, followed by intramembrane cleavage of the C-terminal fragment (CTF) by *γ*-secretase-a process whose structural mechanism has recently been elucidated by cryo-EM (Zhou et al. 2019; Thornton et al. 2017; Feuerbach et al. 2017). This proteolytic sequence effectively terminates TREM2 signalling, enabling microglia to revert to a homeostatic state. Following ligand binding, TREM2 couples with DAP12 via their transmembrane domains (Steiner et al. 2020a; Hamerman et al. 2006; Peng et al. 2010), triggering phosphorylation of DAP12’s immunoreceptor tyrosine-based activation motifs (ITAMs) and recruitment of kinases including SYK, PLC*γ*, and PI3K (Kleinberger et al. 2014; Guerreiro et al. 2013a; Paloneva et al. 2002).

The TREM2-DAP12 interaction is initiated through electrostatic pairing between lysine 186 (K186) of TREM2 and conserved aspartate residues (D50) in DAP12, forming a stable trimeric transmembrane complex (Call et al. 2010a). Additional stabilisation is provided by W194 in TREM2, which forms a hydrogen bond with T54 in DAP12 (Zhong et al. 2024). While most functional studies of TREM2 focus on extracellular domain mutations, emerging evidence implicates transmembrane domain variants in AD risk. Notably, the W191X (X represents truncation) nonsense mutation in TREM2 isoform 2 has been associated with increased AD susceptibility in African American populations (Jin et al. 2015). TREM2 exists in at least three isoforms by alternative splicing (Moutinho et al. 2023; Bharadwaj et al. 2024; Giraldo et al. 2013), with isoform 1 (230 residues) being the canonical and most abundant form (Celarain et al. 2016). Isoform 2 (219 residues), was found to be expressed at lower levels than isoform 1 in the hippocampus of AD patients (Ma et al. 2016), while isoform 3 (222 residues) remains even less studied (Jin et al. 2014).

Structural biology approaches like cryo-EM and X-ray crystallography offer valuable but static views of protein complexes (Callaway 2024; Shoemaker and Ando 2018; Beck et al. 2024). Machine-learning-based prediction models, such as AlphaFold2 and AlphaFold3 (Jumper et al. 2021b; Abramson et al. 2024), provide high-quality structures but often lack the conformational dynamics critical for understanding membrane-embedded interactions (Malhotra et al. 2025). Atomistic molecular dynamics (MD) simulations offer complementary insights by capturing time-resolved structural fluctuations within biologically realistic environments (Karplus and McCammon 2002; Hospital et al. 2015). In this study, we build upon our previous work resolving the TREM2-DAP12 complex in a lipid bilayer (Zhong et al. 2024), investigating the structural consequences of AD-associated and designed point mutations in the TREM2 transmembrane domain (TMD). We investigate seven different forms of TREM2, including four (K186A, K186X, W194A & W194X) mutants of isoform 1 as well as the wild type and two mutants (W191A & W191X) of isoform 2. All simulations are conducted with the TREM2-DAP12 complex embedded within a model lipid membrane consisting of an 80:20 mixture of POPC and cholesterol. By combining multi-scale MD simulations with unsupervised machine learning analysis, we provide an atomistic perspective on how these mutations modulate the TREM2-DAP12 complex stability and their potential implications in Alzheimer’s disease pathogenesis.

## RESULTS

### Unsupervised Learning Reveals Distinct Conformational Clusters in TREM2-DAP12 Complexes

Previous studies have shown that TREM2 interacts with the signalling adaptor DAP12, primarily through a key salt bridge between K186 in TREM2 and D50 in DAP12 (Ulland and Colonna 2018; Steiner et al. 2020b). The cartoons shown in Fig. 1A represent the membrane-embedded *α*-helices structures of TREM2 and DAP12, which we reported in previous work (Zhong et al. 2024). A focused sequence alignment of the transmembrane regions is presented in Fig. 1B (with the full alignment shown in Supplemental Figure S1). We observed that residue W191 in the TREM2-219 isoform corresponds positionally to K186 in TREM2-230. Since K186 is known to be critical for TREM2-DAP12 interaction, and W191 has been potentially linked to Alzheimer’s disease, this suggests that W191 may play a key role in mediating the interaction between TREM2-219 and DAP12.

**Fig. 1.**
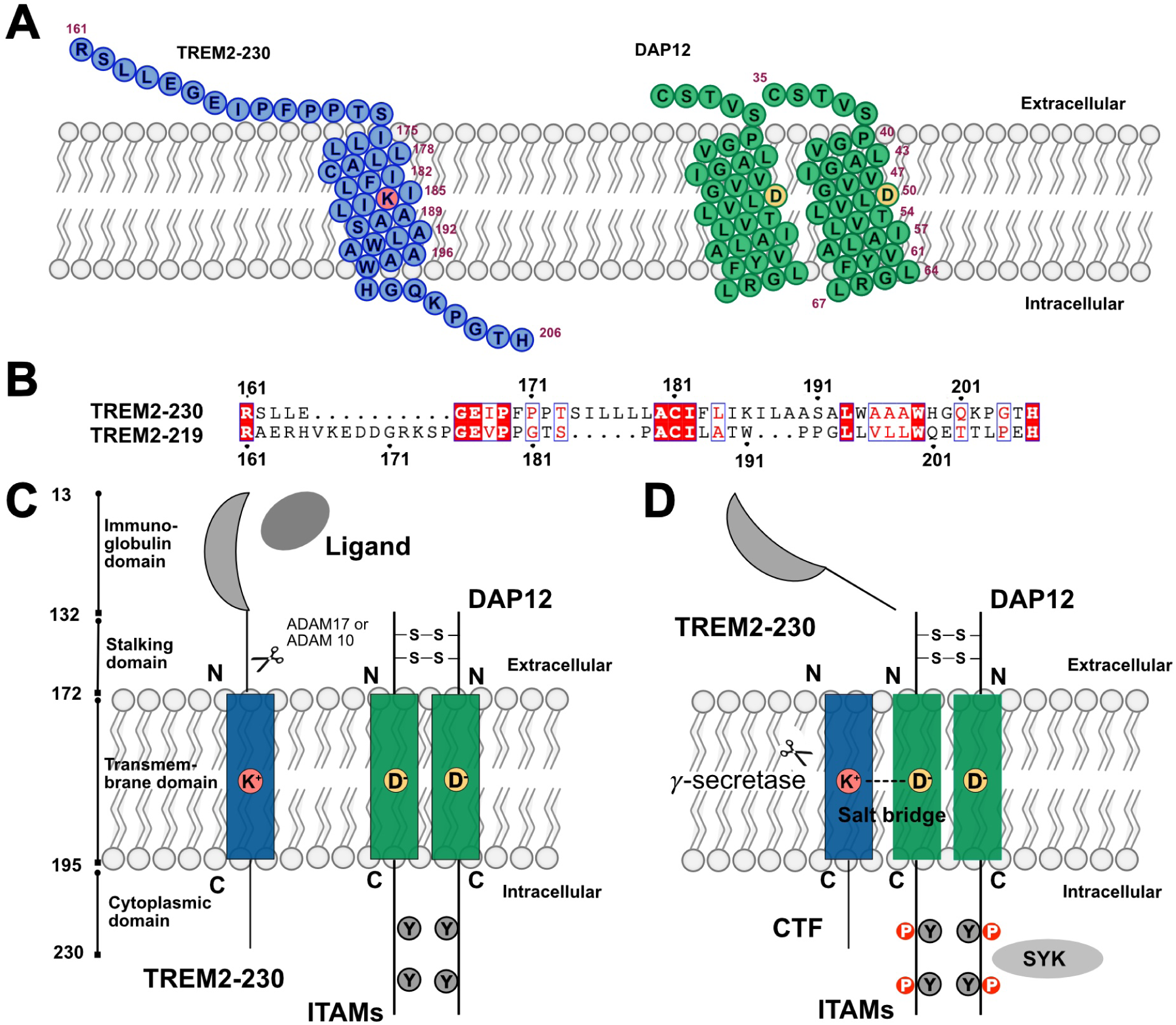
TREM2-DAP12 complex: sequence features and signalling mechanisms. (A) Schematic of the transmembrane domains of TREM2 (isoform 1, 230 aa) and the DAP12 dimer, highlighting key residues involved in membrane embedding and interaction. (B) Sequence alignment of TREM2 isoform 1 (230 aa) and isoform 2 (219 aa), emphasising differences in the transmembrane regions. (C) Ligand engagement induces conformational changes in TREM2, followed by proteolytic cleavage of the ectodomain by ADAM10/17. (D) TREM2-DAP12 association via salt bridges between transmembrane residues triggers downstream phosphorylation of DAP12 ITAM motifs, leading to recruitment of SYK and other kinases.

Fig. 1C illustrates the ligand binding to the extracellular domain of TREM2 and the subsequent intracellular signalling cascade initiated by DAP12. Upon proteolytic cleavage and formation of the C-terminal fragment (CTF), TREM2 remains associated with DAP12, enabling ITAMs phosphorylation and SYK recruitment (Fig. 1D). To investigate how specific mutations affect the structural dynamics of the TREM2-DAP12 complex, we introduced several point mutations into TREM2 and simulated each variant in complex with DAP12. The details of system preparation and mutation design are described in the Methods section. Coarse-grain molecular dynamics (CG-MD) simulations were performed for 300 *μ*s on TREM2-230 variants and 200 *μ*s on TREM2-219 variants.

Using the trajectories from the CG-MD simulations, we implemented an unsupervised machine learning pipeline to characterise the conformational landscapes of the TREM2-DAP12 complexes. High-dimensional contact map data from the trajectories were first embedded into a two-dimensional space using the Uniform Manifold Approximation and Projection (UMAP) algorithm. Then the Hierarchical Density-Based Spatial Clustering of Applications with Noise (HDBSCAN) algorithm was applied to identify distinct conformational states in the UMAP embeddings. The resulting clusters represent dominant structural states sampled by each mutant. These clusters are visualised in Figure 2A and 2D for the TREM2-230 and TREM2-219 constructs, respectively. The relative populations of these conformational states are depicted as doughnut plots in Fig. 2B and Fig. 2E. The temporal evolution of the cluster assignments across the various simulation trajectories, shown in Fig. 2C and Fig. 2F, reveals differences in the stability and dynamics of the sampled conformations of the various mutants.

**Fig. 2.**
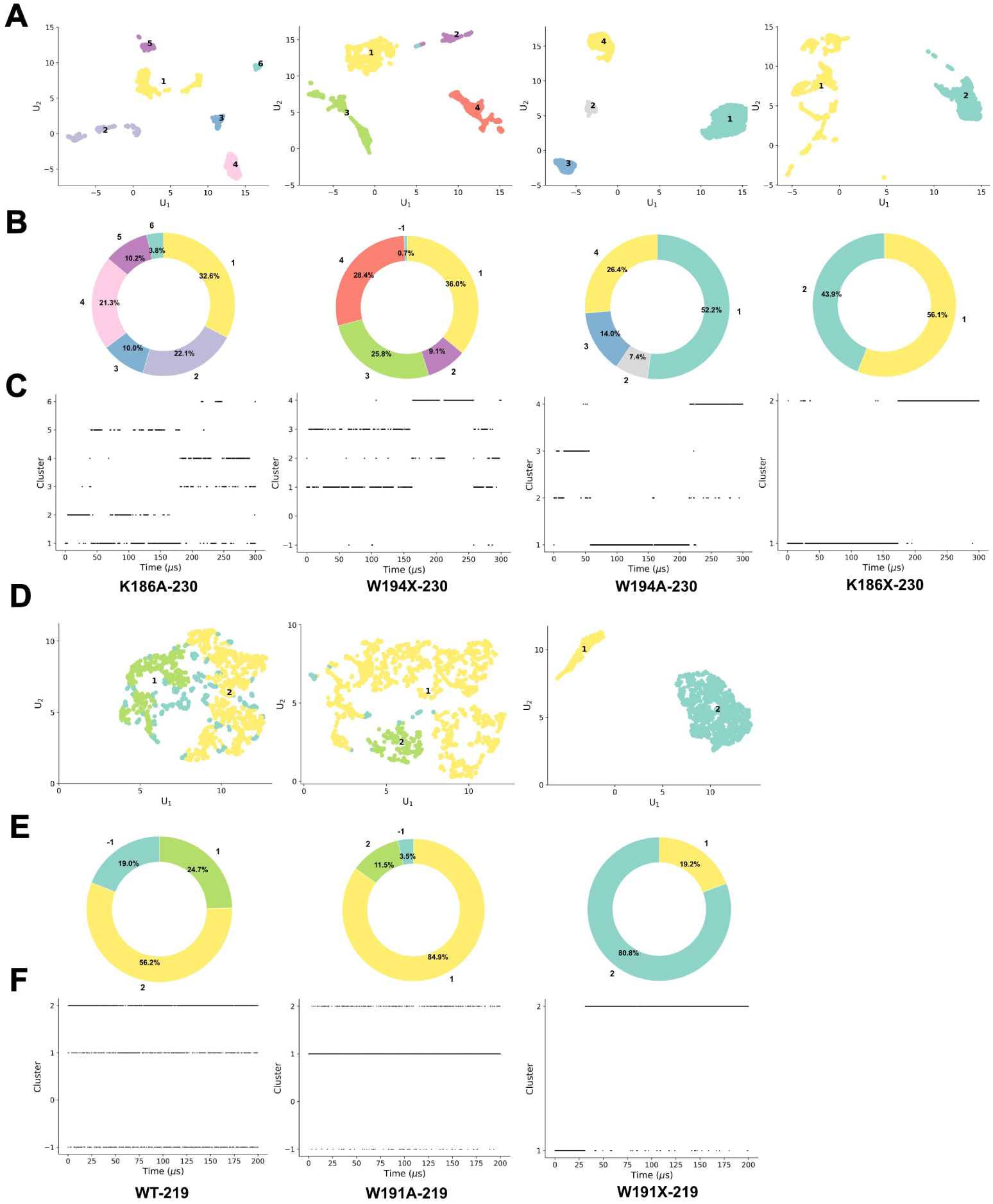
Unsupervised clustering reveals distinct conformational states of TREM2-DAP12 mutants. (A) UMAP (Uniform Manifold Approximation and Projection) plots showing the clustering of conformational states for four TREM2-230 mutants: K186A, W194X, W194A, and K186X. Each point represents a structural snapshot colored by its assigned cluster. (B) Corresponding doughnut charts depicting the relative abundance (percentage) of each conformational cluster identified in panel A. (C) Cluster assignments over time for each mutant in panel A, illustrating the temporal evolution and stability of conformational states across simulation trajectories. (D) UMAP plots of three TREM2-219 constructs: wild-type (WT), W191A, and W191X, showing the distribution of their structural ensembles. (E) Doughnut charts representing the relative population of clusters for the constructs shown in panel D, indicating how mutations alter conformational preferences. (F) Temporal evolution of cluster assignments for the TREM2-219 constructs shown in panel D, highlighting the stability and transitions between conformational states over time.

These results indicate different conformations on different timescales. The detailed contact maps and representative conformational structures are shown in Supplemental Figures S2-S8.

### Characterization of TREM2-DAP12 Interaction Interfaces and Mutant Effects

To further understand the interaction interfaces of the different TREM2-DAP12 complexes, we identified representative structures from the dominant conformational clusters observed in the CG-MD simulations (Table 1) and then converted them to all-atom models using the CHARMM-GUI converter (Qi et al. 2014; Jo et al. 2008). We simulated these resulting systems for 300 ns. From these trajectories, we constructed contact maps using a 3 Å cutoff distance (Zhong et al. 2024), between C-*α* atoms to define residue-residue interactions (Supplemental Figures S9-S15). Residue pairs that maintained contact for over 60% of the trajectory were labelled as interacting residues, while those with contact persistence above 90% were defined as important residues (Fig. 3, blue and red dots, respectively). A comprehensive summary of the interacting residue pairs that persist for 60%, 70%, 80%, and 90% of the trajectories is provided in Supplemental Table S1.

**Fig. 3.**
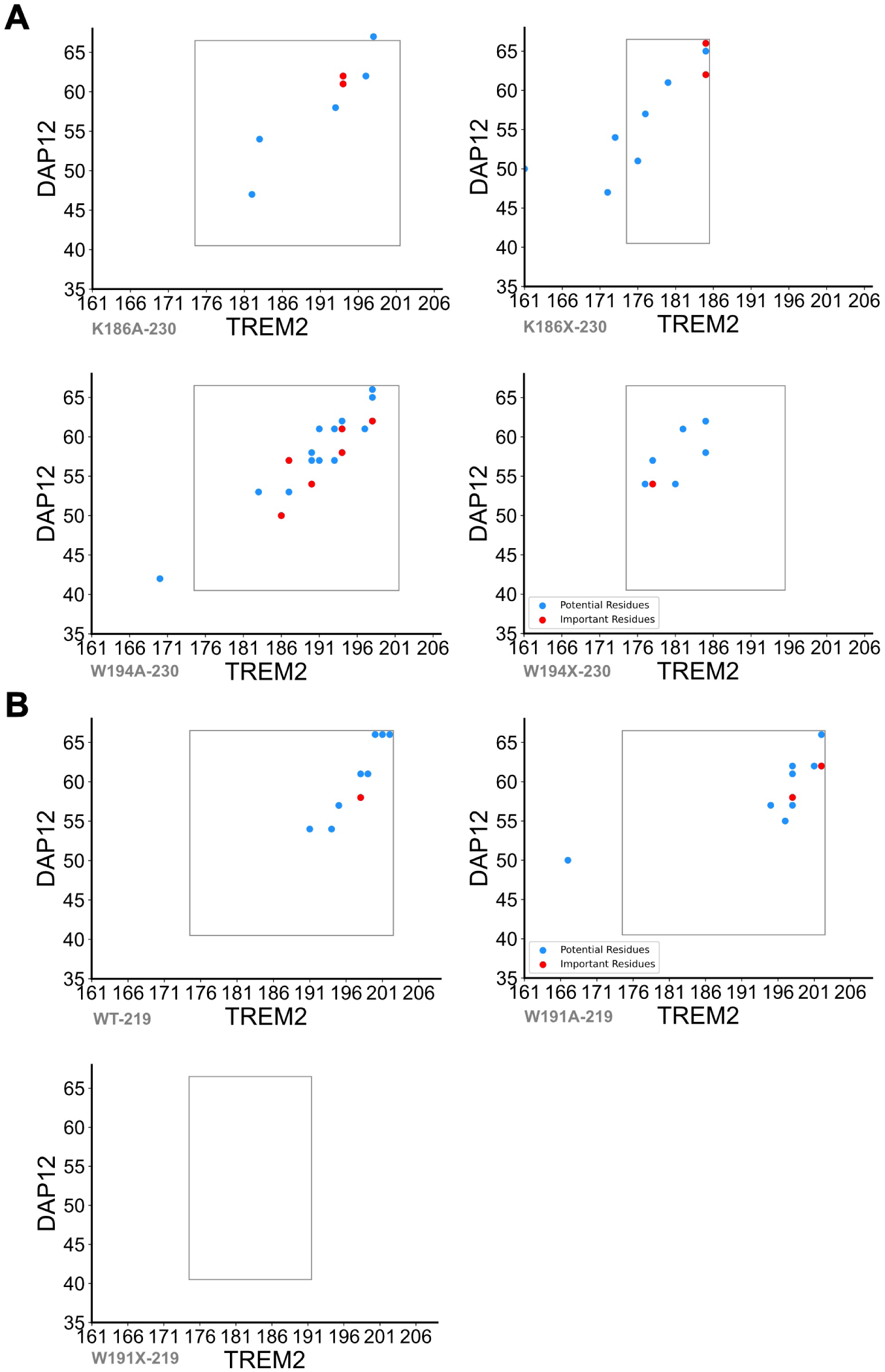
Residue-level interaction analysis of TREM2-DAP12 transmembrane mutants. (A) Pairwise residue interaction maps for TREM2 mutants (K186A, K186X, W191A, W191X) in the 230 construct, plotted with TREM2 residues on the x-axis and DAP12 residues on the y-axis. Blue dots indicate interacting residues, and red dots highlight residues identified as important for the interaction. (B) Residue interaction maps for selected TREM2 constructs in the 219 background: wild-type (WT), W191A, and W191X. These maps reveal the altered interaction profiles resulting from specific point mutations and truncations, underscoring the differential contribution of individual residues to the TREM2-DAP12 interface.

**TABLE 1.** Cluster assignments and times for each variant.

| Variant | Replica 1 | Replica 2 | Replica 3 |
| --- | --- | --- | --- |
| K186A-230 | Cluster 4<br>290 μs | Cluster 1<br>295 μs | Cluster 3<br>298 μs |
| K186X-230 | Cluster 2<br>297 μs | Cluster 4<br>299 μs | Cluster 4<br>300 μs |
| W194A-230 | Cluster 4<br>297 μs | Cluster 4<br>299 μs | Cluster 4<br>300 μs |
| W194X-230 | Cluster 2<br>297 μs | Cluster 2<br>299 μs | Cluster 2<br>300 μs |
| WT-219 | Cluster 1<br>198 μs | Cluster 2<br>199 μs | Cluster 2<br>200 μs |
| W191A-219 | Cluster 2<br>198 μs | Cluster 1<br>199 μs | Cluster 1<br>200 μs |
| W191X-219 | Cluster 2<br>197 μs | Cluster 2<br>198 μs | Cluster 2<br>200 μs |

In isoform TREM2-230 (Fig. 3A), the K186A mutant exhibited fewer persistent contacts than are found in the wild-type, underscoring the role of K186 in stabilising the TREM2-DAP12 complex. The key contact sites between the K186X mutant and DAP12 are found to be in the C-terminal regions of both proteins. Notably, the W194A-230 mutant displayed the highest number of contacts among the isoform 1 variants, with interaction hotspots spanning from the middle to the C-terminal region of TREM2 and across the whole length of DAP12. In contrast, the W194X-230 mutant showed markedly reduced contact frequency, suggesting impaired complex formation.

In isoform TREM2-219 (Fig. 3B), WT-219 complexes retained substantial interactions ranging from the middle to the C-terminal regions of both TREM2 and DAP12. The W191A mutation preserved these contacts with a prominent clustering toward the C-terminal regions. Strikingly, the W191X-219 mutant exhibited no detectable contacts within the defined threshold, indicating a complete loss of interaction. This observation is consistent with the hypothesis that W191 is critical for maintaining the TREM2-DAP12 interface and is in line with genetic studies linking W191 variants to Alzheimer’s disease susceptibility.

The rectangles overlaid in Fig. 3 denote the approximate boundaries of the transmembrane *α*-helices in each complex, providing structural context for the observed contact distributions.

### Mechanistic Insights into Residue-Residue Interactions and Mutant-Induced Disruptions in TREM2-DAP12 Complex Stability

To gain mechanistic insight into the specific residue-residue interactions stabilising TREM2-DAP12 complexes, we analysed interaction types across all mutant systems (Fig. 4A). Within the K186A-230 mutant, we found that *π*-*π* stacking interaction between W194 of TREM2 and Y62 of DAP12 was observed in the K186A-230 mutant (Fig. 4B) to play an important in the formation of the complex, occurring 37.60% of the total simulation time-32.28% in a parallel (sandwich-like) orientation and 5.32% in a T-shaped configuration (Fig. 4C). The details of the *π*-*π* stacking measurement can be found in Supplemental Figure S16. In addition, a hydrophobic interaction between W194 and V61 further supports trimer stability (Fig. 5A).

**Fig. 4.**
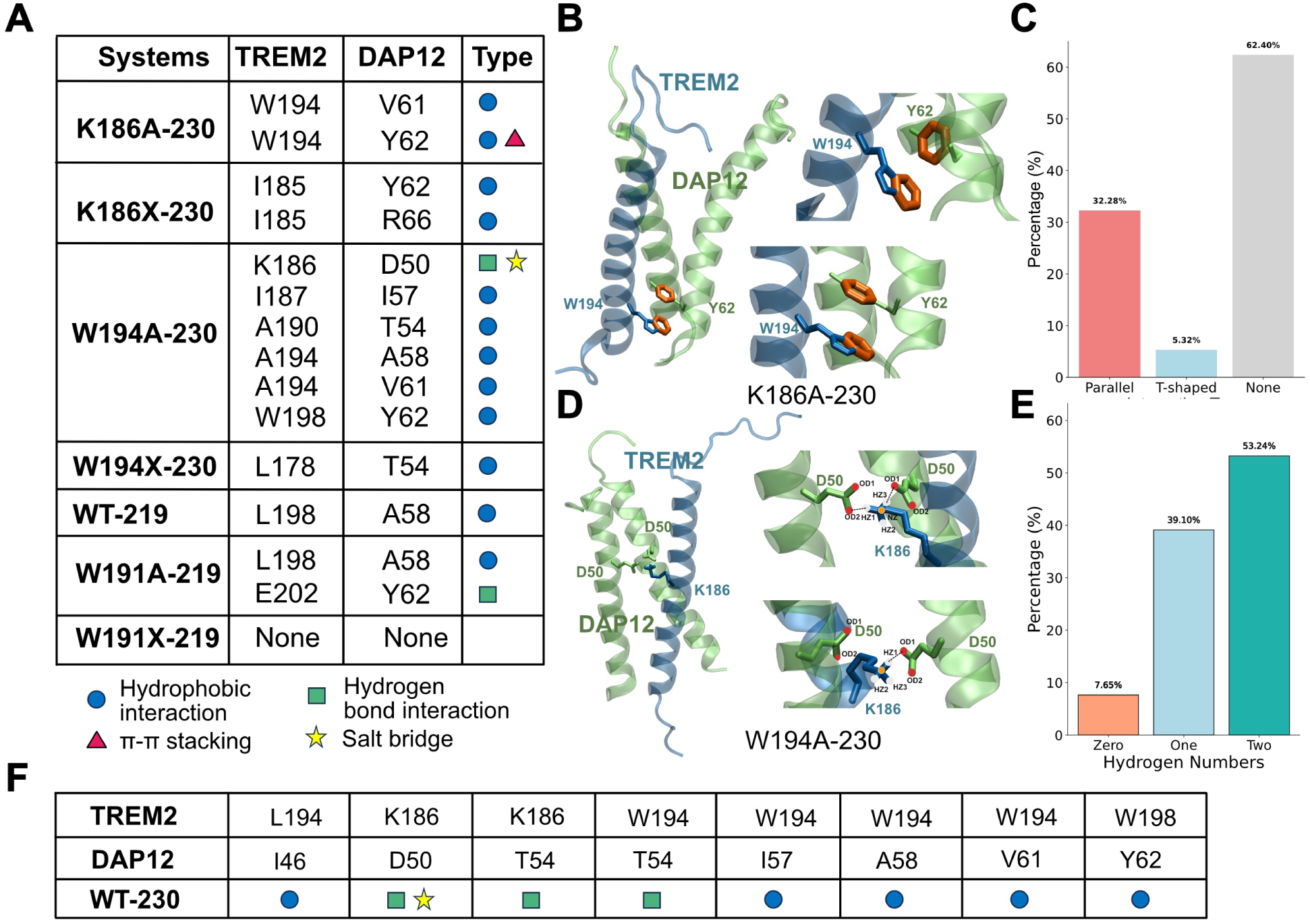
Molecular interaction profiles of TREM2-DAP12 transmembrane mutants. (A) Table summarising key residue-residue interactions between TREM2 and DAP12 for various mutants and constructs. Interaction types are colour-coded: hydrophobic (blue circle), *π*-*π* stacking (red triangle), hydrogen bonds (green square), and salt bridges (yellow star). (B) Representative structural models illustrating *π*-*π* stacking interactions between W194 (TREM2) and Y62 (DAP12) observed in K186A-230. (C) Bar graph quantifying the frequency of different *π*-*π* interaction geometries (parallel, T-shaped, or none) across simulation frames. (D) Structural snapshots showing the formation of a salt bridge between K186 (TREM2) and D50 (DAP12) in W194A-230. (E) Histogram showing the distribution of hydrogen bond numbers formed between K186 and D50 across simulations. These analyses demonstrate that specific point mutations significantly alter interaction patterns at the TREM2-DAP12 interface. (F) The Table shows the important residue pairs of WT-230.

**Fig. 5.**
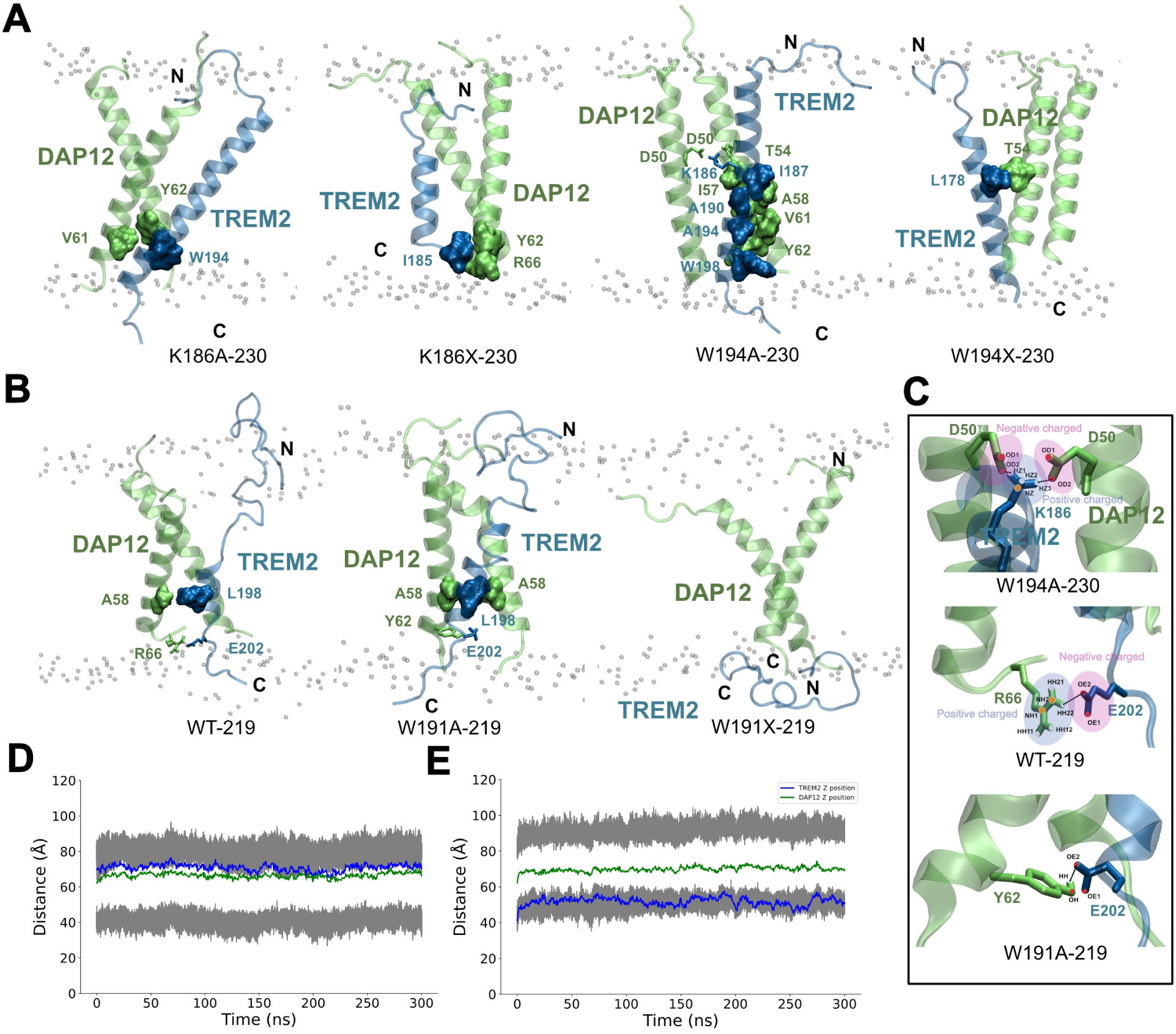
Structural insights into the TREM2-DAP12 transmembrane interactions and stability across mutants. (A) Representative snapshots of the TREM2-DAP12 transmembrane complexes embedded in a lipid bilayer for the TREM2-230 mutants: K186A, K186X, W194A, and W194X. Interacting residues at the interface are shown in surface representation. (B) Structural models of TREM2-219 constructs (WT, W191A, W191X) in complex with DAP12, highlighting the critical interfacial residues. (C) Zoomed-in views of key electrostatic and hydrogen-bond interactions formed between TREM2 and DAP12 residues in W194A-230, WT-219, and W191A-219. Positively and negatively charged regions are indicated. (D, E) Distance measurements over time between the centre of mass of selected TREM2 and DAP12 transmembrane residues from molecular dynamics simulations. (D) shows the distance with the WT-219; (E) shows constructs with the W191X-219 construct. These structural and dynamic analyses reveal that specific point mutations and truncations significantly alter the interfacial interactions and complex stability.

In the W194A-230 mutant, a stable hydrogen bond forms between K186 in TREM2 and D50 in both DAP12 monomers (Fig. 4D). This hydrogen bond persists for 92.34% of the simulation, with a hydrogen bond to one DAP12 monomer present 39.10% of the time and hydrogen bonds to both DAP12 monomers occurring simultaneously in 53.24% of the frames (Fig. 4E). The details of the number of hydrogen bonds as a function of time can be found in Supplemental Figure S17. The further interaction network that stabilises the W194A-230 complex includes multiple hydrophobic contacts and a salt bridge that contribute to tight trimer packing (Fig. 5C, top panel). To facilitate comparison between the wild-type and mutant forms of isoform 1, the previously reported key interacting residues in WT-230 (Zhong et al. 2024) are also highlighted in Fig. 4F.

In contrast, the two truncated TREM2-230 complexes are primarily stabilised by hydrophobic interactions. In the case of the W194X-230 complex, the hydrophobic contact between L178 (TREM2) and T54 (DAP12) is found to play an important role in the stabilisation of the complex. While in the K186X-230 complex, we found that the complex is a relatively stable conformation driven by hydrophobic interactions involving I185 of TREM2 with Y62 and R66 of DAP12 (Fig. 5A).

For the WT-219 system, complex stability is maintained through a hydrophobic interaction between L198 (TREM2) and A58 (DAP12), along with a salt bridge and hydrogen bond between E202 (TREM2) and R66 (DAP12), with a 70% occupancy reported in Table S1 and shown in Fig. 5C (middle panel). The W191A-219 mutant retains the L198-A58 hydrophobic interaction and features an additional hydrogen bond between E202 (TREM2) and Y62 (DAP12) (Fig. 5C, bottom panel).

In stark contrast, the W191X-219 mutant completely disrupts all residue-level contacts, decreasing complex stability (Fig. 4A). Structural analysis reveals that TREM2 reorients horizontally along the membrane interface, while DAP12 retains a vertical orientation (Fig. 5B). Analysis of the *z*-axis positioning over time (Fig. 5D and Fig. 5E) further confirms this dissociation: in WT-219, both proteins remain colocalised in the upper membrane leaflet, whereas in W191X-219, TREM2 shifts toward the lower leaflet while DAP12 remains within the center of the membrane.

The K186X-230 and W191X-219 mutants exhibited significantly greater center-of-mass distances between TREM2 and DAP12 in the *x* − *y* plane, measuring 13.02 Å and 11.20 Å, respectively (Supplementary Figure S18). In contrast, smaller distances were observed in the K186A-230 (7.86 Å), W194A-230 (8.08 Å), W194X-230 (9.83 Å), WT-219 (6.06 Å), and W191A-230 (6.07 Å) complexes. These findings suggest that the increased spatial separation in K186X-230 and W191X-230 reflects destabilised or weakened interactions between TREM2 and DAP12.

### Impact of Point Mutations on Structural Flexibility and Stability of the TREM2-DAP12 Complex

To investigate how point mutations affect structural flexibility, we analysed the root mean square fluctuation (RMSF) profiles of each TREM2-DAP12 complex (Fig. 6). Fig. 6 A shows RMSF values for the 230-residue constructs. Compared to the WT-230 complex, the K186A-230 mutant displays increased flexibility in the N-terminal region of TREM2. This likely results from the loss of K186-mediated anchoring interactions. In the K186X-230 complex, higher fluctuations are observed in one of the DAP12 chains, suggesting that truncation of TREM2 disrupts the symmetric stability of the trimeric interface. The W194A-230 variant also exhibits elevated dynamics in the N-terminal domain of TREM2, possibly due to the loss of the aromatic W194 residue, which contributes to *π*-stacking and hydrophobic stabilisation. W194X-230 shows localised fluctuation increases but remains relatively constrained compared to K186X.

**Fig. 6.**
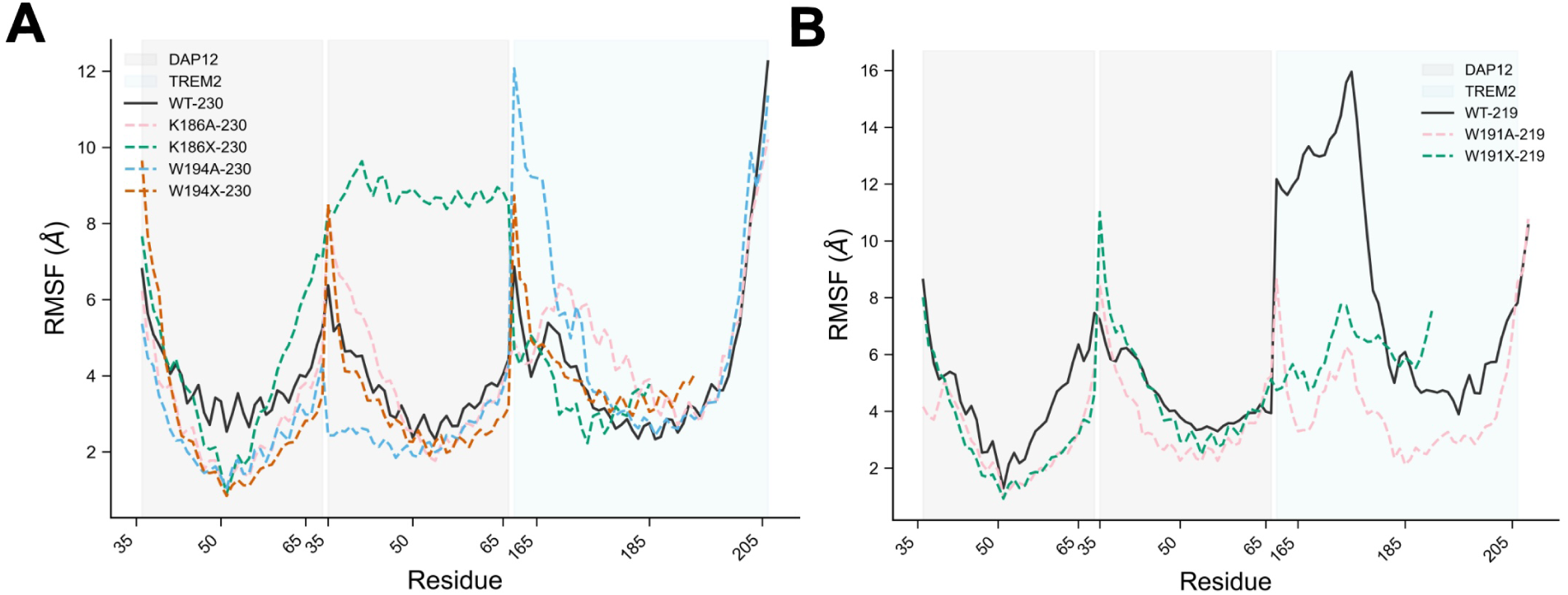
Root Mean Square Fluctuation (RMSF) profiles of TREM2-DAP12 complexes highlight dynamic changes due to mutations. (A) RMSF plots of residue-wise flexibility for DAP12 and TREM2 in the 230-residue constructs. WT-230 and four mutants (K186A, K186X, W194A, W194X) are compared. Shaded regions denote the DAP12 (gray) and TREM2 (blue) chains. Mutations lead to distinct increases in flexibility, particularly in the transmembrane and cytoplasmic regions. (B) RMSF profiles for the 219-residue constructs: WT-219, W191A-219, and W191X-219.

In the 219-residue systems (Fig. 6B), the WT-219 complex exhibits extensive flexibility in the cytoplasmic region of TREM2 and the DAP12 interface, maybe due to the lack of interaction of the N-terminus of TREM2 with either DAP12 or lipids (Fig. 5B). W191A-219 displays a slightly more rigid profile, possibly due to compensatory stabilising hydrogen bonds. W191X-219 shows moderate flexibility across both proteins. These RMSF profiles underscore the dynamic nature of the TREM2-DAP12 interaction and reveal how specific mutations perturb stability through altered flexibility in both transmembrane and cytoplasmic domains.

## DISCUSSION

The W191X mutation introduces a premature stop codon within the transmembrane domain of the TREM2-219 isoform, leading to truncation of the receptor’s C-terminal tail. Genetic studies have previously associated this nonsense variant with an increased risk of Alzheimer’s disease, particularly in African American populations (Giraldo et al. 2013; Jin et al. 2015). However, the biophysical consequences of W191X on TREM2-DAP12 complex formation and function have remained elusive.

Our simulations provide structural and mechanistic insights into how this mutation disrupts interprotein signalling. In contrast to the WT-219 and W191A-219 systems, which maintain robust interactions through hydrophobic, hydrogen bond, and salt bridge networks, the W191X-219 complex shows a complete loss of residue-level contacts at the TREM2-DAP12 interface (Fig. 3B & Fig. 4A). This disruption is further supported by the W191X variant, which exhibits a striking decoupling of the two helices, both laterally (11.2 Å in the *x*-*y* plane) and vertically (along the *z*-axis) orientation (Fig. 5D, Fig. 5E & Supplemental Figure S18). Notably, the TREM2 helix in W191X-219 reorients almost horizontally along the bilayer interface-a configuration incompatible with canonical receptor-adaptor signalling (Colonna 2023; Ulland and Colonna 2018).

At the molecular level, the truncation eliminates distal cytoplasmic residues, including E202-a key site for salt bridge formation with R66 of DAP12 in WT-219 and W191A-219 systems (Fig. 5C). The loss of this interaction likely destabilises the helix alignment necessary for signal transduction via ITAMs on DAP12. These findings collectively suggest that W191X disrupts not only static contacts but also the dynamic coherence required for functional signalling.

Importantly, our results also provide comparative insights into mutation-induced destabilisation across isoforms. The K186X-230 mutant, though distinct in sequence and location, mirrors the disruption seen in W191X-219, with increased centre-of-mass separation and reduced interhelical stability (Fig. 5A, Supplemental Figure S18). This parallel underscores the critical role of C-terminal residues in preserving the spatial geometry of the receptor-adaptor complex.

Beyond the immediate biophysical implications, these findings have broader relevance for understanding TREM2’s role in neuroimmune signalling. Misalignment or dissociation of TREM2 from DAP12 could impair SYK recruitment and ITAMs phosphorylation, mechanisms essential for microglial activation and phagocytosis (Wang et al. 2022; Zhang et al. 2025). Given TREM2’s critical role in regulating microglial responses to amyloid plaques, including early-stage plaque seeding and clustering (Parhizkar et al. 2019; Ulrich et al. 2017), structural disruptions such as W191X may mechanistically link altered microglial function to increased Alzheimer’s disease risk.

In summary, this study establishes a molecular framework for understanding how C-terminal truncation mutations, such as W191X, destabilise the TREM2-DAP12 complex. It underscores the importance of structural integrity in maintaining signalling-competent receptor assemblies and provides new insights for targeting TREM2 variants in the context of neurodegenerative diseases.

## METHODS

### Approach to Build Systems

The TREM2 mutants are made with ColabFold v1.5.2 (Mirdita et al. 2022; Jumper et al. 2021a) using the mutated sequence (structures confidence can be found in Supplemental Figure S19-S25), and the DAP12 dimer is using PDB 2L34 (Call et al. 2010b). The complexes are made by merging the TREM2 mutants and DAP12 together as a reference using HADDOCK (Dominguez et al. 2003; Van Zundert et al. 2016), AlphaFold2 (Jumper et al. 2021a), AlphaFold3 (Abramson et al. 2024), and the Wild-type complex structure, we simulated a stable conformation. The sequences of each system can be found in Supplemental Table S2. Each system was then equilibrated using all-atom molecular dynamics simulations for 100 ns. The systems utilised in this paper were generated through the use of the CHARMM-GUI Membrane Builder (Jo et al. 2008) and the MARTINI Maker (Siewert et al. 2007) for all-atom (AA) and coarse-grain (CG) simulations, respectively. The membrane consists of POPC and cholesterol at a ratio of 80 : 20. The solvent employed in every simulation was water, and to maintain a neutral charge and a 0.15 M salt concentration, Na^+^ and Cl^−^ ions were added to the system. The initial configurations for each system are shown in Supplemental Figure S26, while the complete simulation workflow is outlined in Supplemental Figure S27.

### Coarse-grain Molecular Dynamics Simulations

Coarse-grain simulations of 300 *μ*s and 200 *μ*s were performed for isoform 1 and isoform 2, respectively, to obtain equilibrated systems embedded in a POPC/cholesterol lipid bilayer. Simulations were carried out using GROMACS version 2019.2 (Van Der Spoel et al. 2005). The MARTINI22P force field (Qi et al. 2015) was applied for all systems. Each membrane system underwent energy minimisation followed by equilibration at 310.15 K. The MARTINI22P model maps three to five non-hydrogen atoms onto coarse-grain beads, with bead types defined by the atomic composition and chemical characteristics. Beads are categorised into four main classes-charged, polar, non-polar, and apolar-with five subtypes per class, distinguished by polarity and hydrogen-bonding potential. An overview of the CG simulation setups for various TREM2-DAP12 complexes is provided in Table 2.

**TABLE 2.** Details of each of the coarse-grain MD simulation systems.

| **Variant** | $n_{\text{beads}}$ | $n_{\text{lipids}}$ | $n_{\text{waters}}$ | $n_{\text{ions}}$ | **Final box dimensions (nm)** |
| --- | --- | --- | --- | --- | --- |
| K186A-230 | 24024 | 310 | 6713 | 152 | $100.6 \times 100.6 \times 127.4$ |
| K186X-230 | 19022 | 310 | 5075 | 115 | $100.2 \times 100.2 \times 107.0$ |
| W194A-230 | 23904 | 310 | 6673 | 153 | $100.4 \times 100.4 \times 127.2$ |
| W194X-230 | 19618 | 310 | 5267 | 118 | $100.3 \times 100.3 \times 108.7$ |
| WT-219 | 24875 | 310 | 6089 | 158 | $100.5 \times 100.5 \times 130.6$ |
| W191A-219 | 18802 | 310 | 4980 | 116 | $100.5 \times 100.5 \times 106.1$ |
| W191X-219 | 24558 | 320 | 6883 | 154 | $100.7 \times 100.7 \times 128.7$ |

### All-atom Molecular Dynamics Simulations

Following 300 *μ*s or 200 *μ*s of coarse-grain simulations, each system was converted to an all-atom representation using the CHARMM-GUI Martini to All-atom Converter (Qi et al. 2014; Jo et al. 2008), and subsequently simulated for 300 ns. GROMACS version 2020.3 (Van Der Spoel et al. 2005) was used to perform these AA simulations. The same POPC/cholesterol bilayer composition was retained, with lipids described by the CHARMM36 force field (Guvench et al. 2011; Klauda et al. 2010) and water represented by the CHARMM-modified TIP3P model (MacKerell Jr et al. 1998). Each system was first energy-minimised and then equilibrated at 310.15 K and 1 bar, following CHARMM-GUI protocols (MacKerell Jr et al. 1998; Best et al. 2012). The equilibration began with steepest descent minimisation, followed by a 125 ps NVT ensemble run using the Nosè-Hoover thermostat to maintain temperature (310.15 K) with a 1 fs time step. This was followed by a 300 ns production simulation in the NPT ensemble, employing the Nosè-Hoover thermostat and the Parrinello-Rahman barostat to sustain the temperature at 310.15 K and pressure at 1 atm (NOSÉ 2002; Parrinello and Rahman 1981). Bond constraints involving hydrogens were applied using the LINCS algorithm (Hess et al. 1997). Periodic boundary conditions were used in all three dimensions. Both equilibration and production phases used a 1 fs time step. Electrostatic and Lennard-Jones (LJ) interactions were truncated at 12 Å, with a switching function applied to taper LJ interactions to zero starting from 10 Å. Table 3 summarises the AA simulation setups for the TREM2-DAP12 complexes. Three independent replicas were conducted.

**TABLE 3.** Details of each of the all-atom MD simulation systems.

| **Variant** | $n_{\text{atoms}}$ | $n_{\text{lipids}}$ | $n_{\text{waters}}$ | $n_{\text{ions}}$ | **Final box dimensions (nm)** |
| --- | --- | --- | --- | --- | --- |
| K186A-230 | 120584 | 310 | 26969 | 152 | $97.9 \times 97.9 \times 130.1$ |
| K186X-230 | 100554 | 310 | 20405 | 115 | $97.7 \times 97.7 \times 106.9$ |
| W194A-230 | 120055 | 310 | 26793 | 153 | $97.4 \times 97.4 \times 130.7$ |
| W194X-230 | 103026 | 310 | 21188 | 118 | $96.1 \times 96.1 \times 113.5$ |
| WT-219 | 123893 | 310 | 28060 | 158 | $98.3 \times 98.3 \times 132.0$ |
| W191A-219 | 99717 | 310 | 20020 | 116 | $99.2 \times 99.2 \times 103.7$ |
| W191X-219 | 122945 | 315 | 27642 | 154 | $98.5 \times 98.5 \times 130.3$ |

### Simulation Analysis Techniques

All simulations were analysed using in-house Python (3.10) scripts built primarily on the MDAnalysis package (Gowers et al. 2016; Michaud-Agrawal et al. 2011). Visualisations were generated using Matplotlib (Hunter 2007), and trajectory inspection was carried out with VMD 1.9.3 (Humphrey et al. 1996).

#### Hydrogen Bond Analysis

Hydrogen bond probabilities were calculated using the Hydrogen Bond Analysis module in MDAnalysis (Smith et al. 2019), via custom Python scripts. A donor-acceptor distance cutoff of 3 Å and a donor-hydrogen-acceptor angle cutoff of 150 ^◦^ were applied.

#### *π*-*π* Stacking Analysis

*π*-*π* stacking interactions were evaluated using an in-house Python script based on PySoftK (López-Ríos de Castro et al. 2025). Ring centroids were computed as the mean position of ring atoms, and ring plane orientation was defined using three non-collinear atoms. Interactions were classified using a centroid distance cutoff of 6.5 Å (Piovesan et al. 2016) and an interplanar angle criterion: ≤30^◦^ for parallel-displaced, and 60^◦^-120^◦^ for T-shaped stacking.

#### Conformational Clustering of TREM2-DAP12 Complexes

To identify stable conformational states of the TREM2-DAP12 complexes from coarse-grain simulations, we first extracted inter-residue distance features from backbone beads (BB) spaced every five residues along the transmembrane helices of TREM2 and both DAP12 chains. At each simulation frame, we computed pairwise distance matrices between the selected BB atoms of TREM2 and those of each DAP12 chain independently using MDAnalysis. These matrices were flattened into feature vectors and concatenated across frames to form a high-dimensional dataset representing the dynamic spatial organisation of the transmembrane helices.

We then applied UMAP for dimensionality reduction, followed by clustering with HDBSCAN. These hyperparameters were selected iteratively to yield physically meaningful conformational clusters. The resulting low-dimensional embeddings captured dominant structural states sampled by each system and were used to initialise subsequent all-atom simulations. Previously, we have used this general approach to characterising the conformations taken by antimicrobial peptides (Serian et al. 2024), apo-c3 (al Badri et al. 2022), interacting lipids (Smith et al. 2020) and polymers (Ziolek et al. 2021; Ziolek et al. 2022; López-Ríos de Castro et al. 2023; López-Ríos de Castro et al. 2024). For isoform 1, UMAP was performed with n_neighbors set to 50, and subsequent clustering used HDBSCAN with min_cluster_size = 50 and cluster_selection_epsilon = 2.0. For isoform 2, the same min_cluster_size was used, with n_neighbors = 10 and cluster_selection_epsilon = 1.5.

#### Contact Map Analysis

Residue-level contact maps between TREM2 and DAP12 were calculated using a custom Python script with MDAnalysis. Contacts were defined as any pair of heavy atoms from TREM2 and DAP12 residues within 3 Å. At each frame, a binary contact matrix was generated and accumulated over the trajectory. The final contact map represents the frequency of residue-residue contacts over time, enabling identification of stable interaction interfaces.

#### Root Mean Square Fluctuation (RMSF) Analysis

RMSF was calculated using a custom Python script with MDAnalysis and NumPy. Alpha carbon atoms were selected, and the trajectory was iterated to compute the mean atomic positions and displacement over time. The RMSF for each residue was computed as:

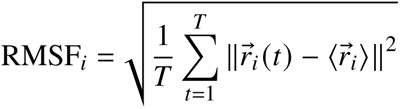

where 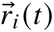 is the position of atom *i* at frame *t*, and 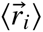 is its time-averaged position. This analysis was performed across the full trajectory for each system to assess per-residue flexibility.

## ACKNOWLEDGEMENT

We thank Dr. Robert M. Ziolek for his valuable feedback on our manuscript. We are grateful to the UK Materials and Molecular Modelling Hub for computational resources, which is partially funded by EPSRC (EP/T022213/1, EP/W032260/1 and EP/P020194/1). We also appreciate the King’s Computational Research, Engineering and Technology Environment (CREATE) at King’s College London for providing us access to computational resources (e Research 2025). Z.Z acknowledges receiving PhD studentships from King’s College London and China Scholarship Council.

## CONTRIBUTION

Z.Z, M.B.U, and C.D.L designed research; Z.Z performed research; Z.Z and C.D.L analysed data; Z.Z and C.D.L wrote the paper.

## DATA AVAILABILITY

Example trajectories and system setup for analysis are available in a Zenodo repository, accessible at https://doi.org/10.5281/zenodo.15746936.

## SUPPLEMENTARY INFORMATION

Supplementary Figures and Tables are available as a PDF.

