## Supplemental Figures and Tables for "Structural Impact of Alzheimer’s Disease Mutations on Transmembrane TREM2-DAP12 Interactions: An Atomistic Perspective"

### Supporting Information for "Structural Impact of Alzheimer's Disease Mutations on Transmembrane TREM2-DAP12 Interactions: An Atomistic Perspective"

Zhiwen Zhong,[†,‡] Martin B. Ulmschneider,[‡] and Christian D. Lorenz[*,¶]

†*Department of Physics, King's College London, London, WC2R 2LS, UK*

‡*Department of Chemistry, King's College London, London, SE1 1DB, UK*

¶*Department of Engineering, King'sCollege London, London WC2R 2LS, UK*

**This PDF file includes:**

Figure S1 to S27

Tables S1 to S2

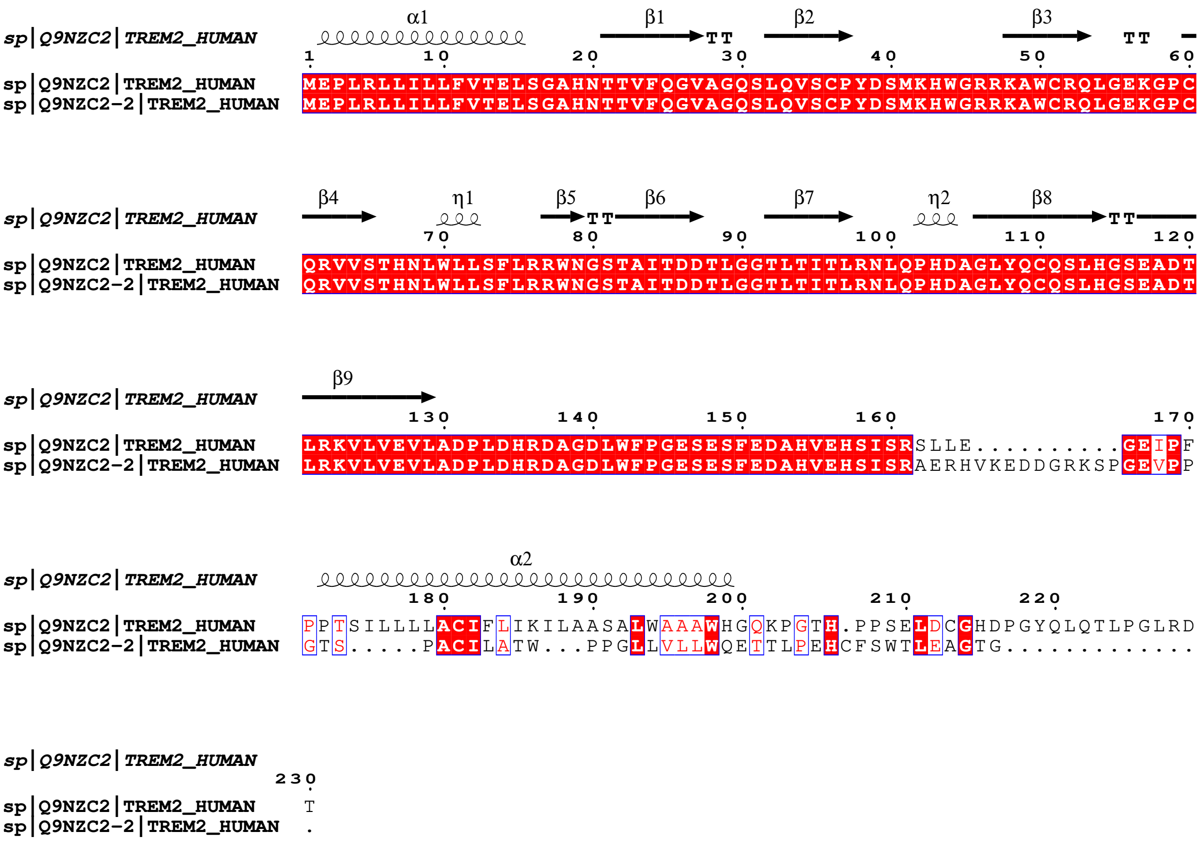


Figure S1: Sequence alignment of two TREM2 isoforms. Isoform 1 (Q9NZC2) corresponds to the full-length 230-amino-acid sequence, while isoform 2 (Q9NZC2-2) is a truncated variant consisting of 219 residues. Secondary structure annotations for isoform 1 were predicted using AlphaFold2 and are indicated above the sequences. The alignment highlights the conserved regions and structural differences between the two isoforms.

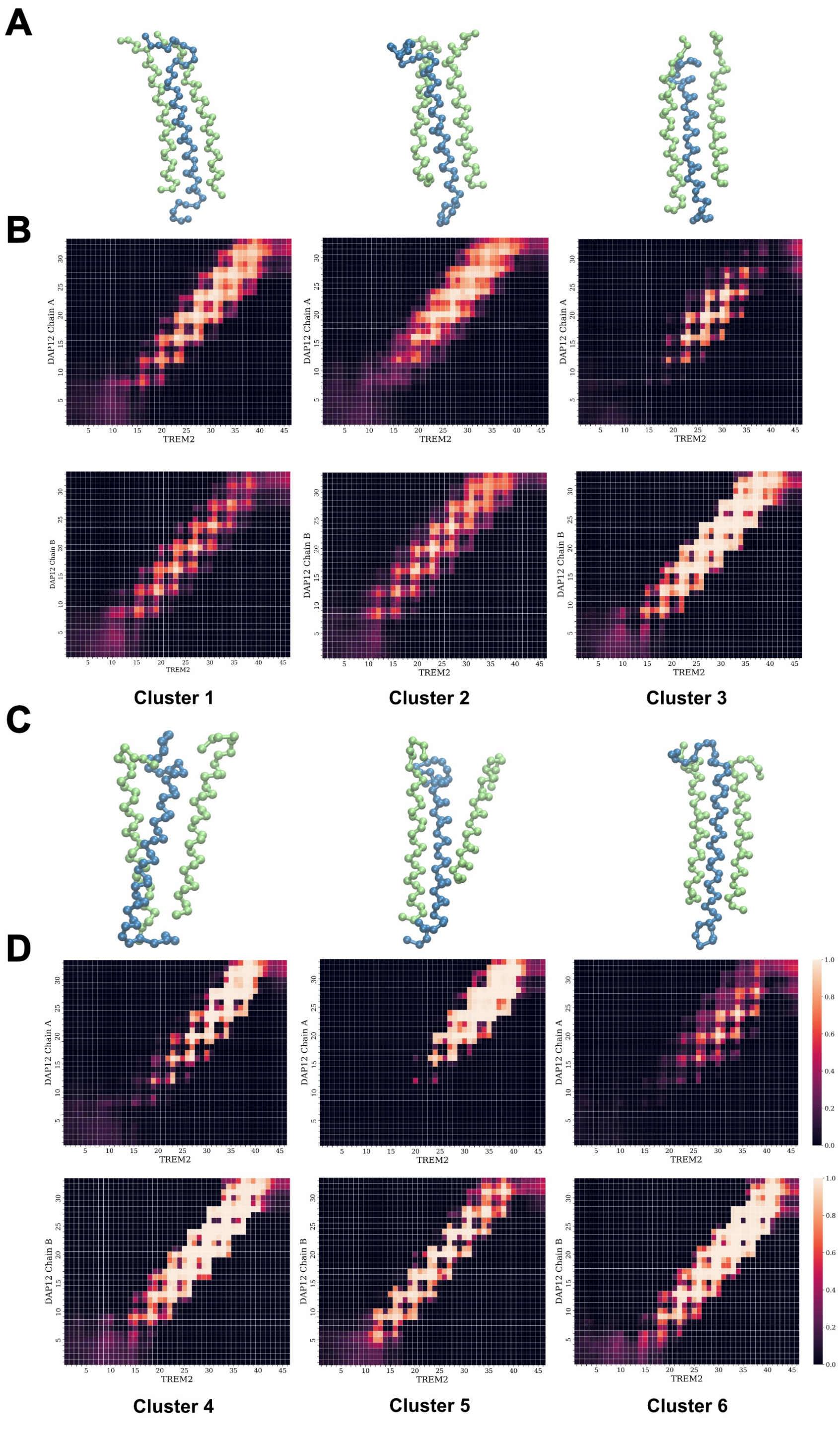


Figure S2: Structural ensembles and contact frequency maps of the K186A-230 system. (A) Representative transmembrane conformations of Cluster 1, Cluster 2, and Cluster 3 from the K186A-230 system, extracted from the clusters based on distance-based clustering of MD trajectories. (B) Corresponding contact maps showing contacts between TREM2 and DAP12 Chain A and Chain B for Clusters 1-3. Contact frequency is color-coded, with lighter shades indicating higher persistence. (C) Representative transmembrane conformations of Cluster 4, Cluster 5, and Cluster 6. (D) Contact frequency matrices for Clusters 4-6, presented as in panel B. The matrices reflect stable inter-residue interactions that define the unique packing arrangements of each cluster.

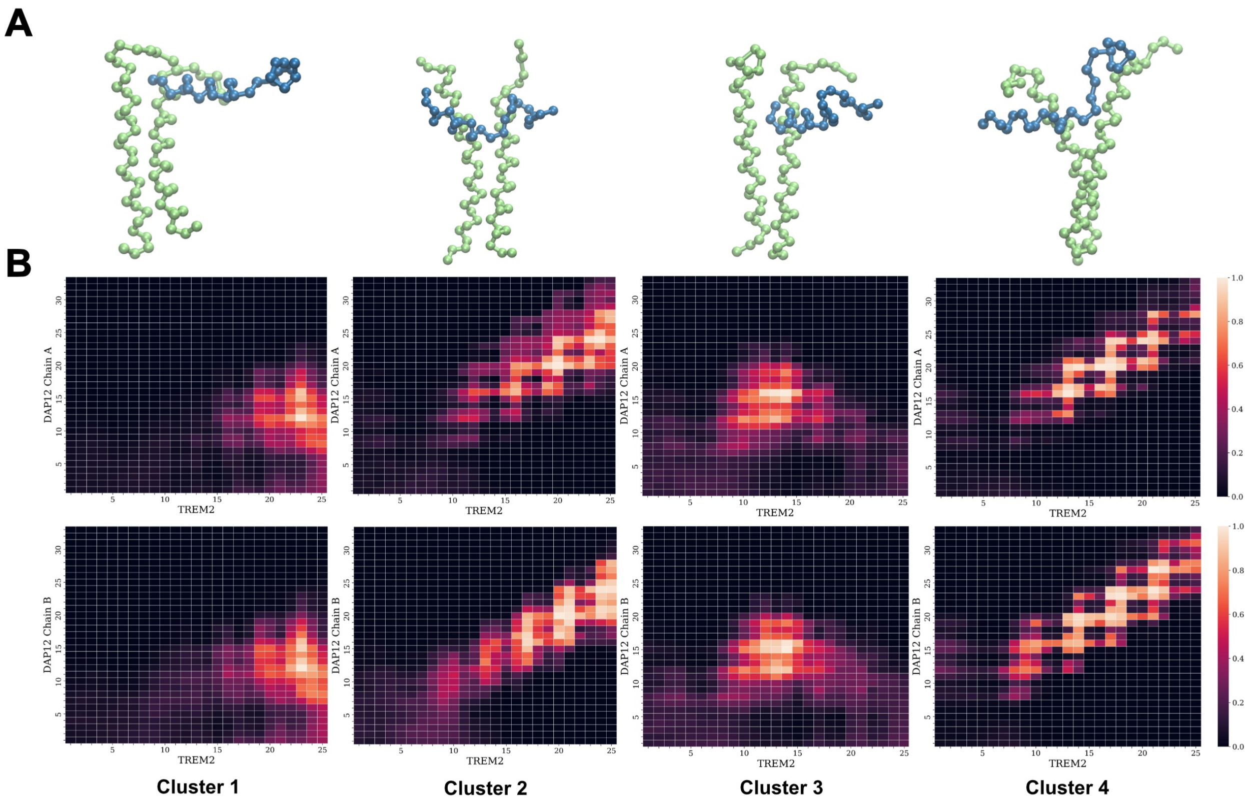


Figure S3: Structural ensembles and contact frequency maps of the K186X-230 system. (A) Representative transmembrane conformations of Cluster 1 to Cluster 4, derived from distance-based clustering of MD trajectories. The structural models illustrate diverse orientations and packing arrangements of the TREM2-DAP12 complex in the absence of K186. (B) Corresponding contact frequency maps between TREM2 and DAP12 Chain A and Chain B for Clusters 1-4. Contact frequency is colour-coded from low (dark) to high (light), highlighting distinct interaction patterns that characterise each conformational state.

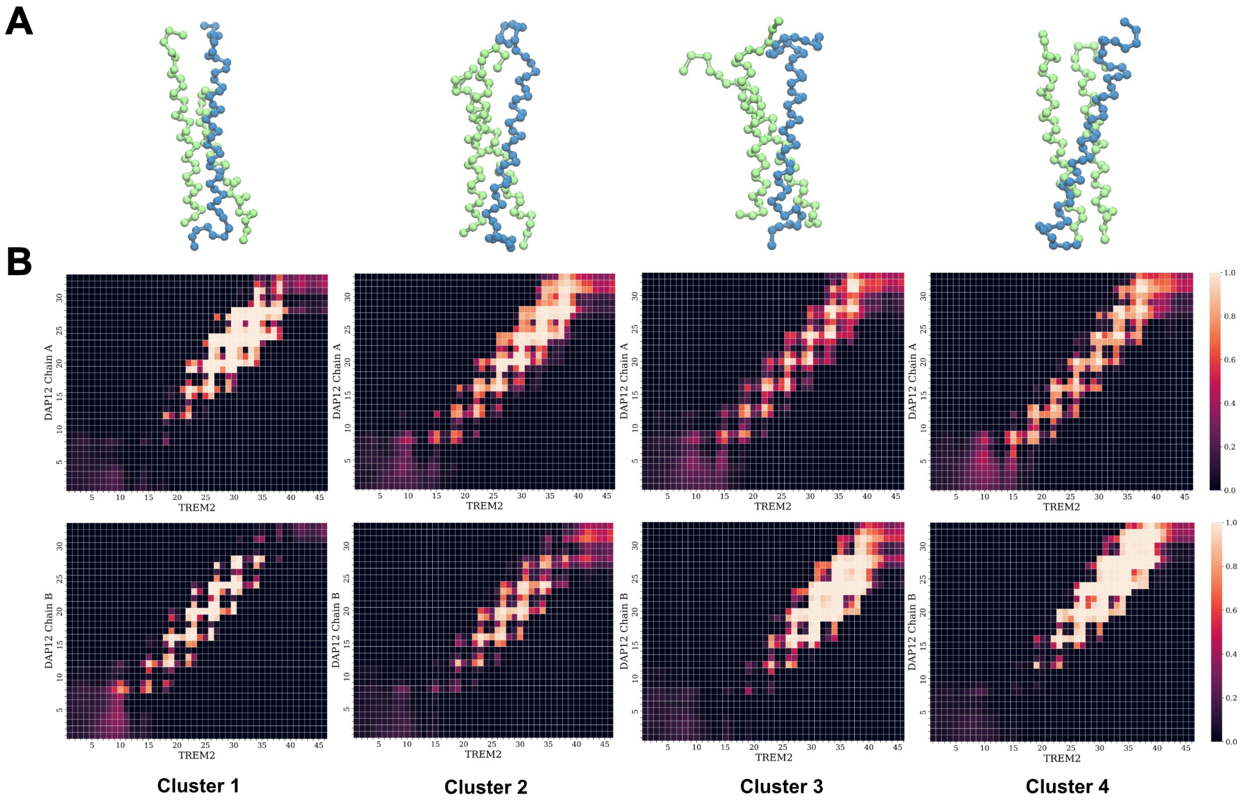


Figure S4: Structural ensembles and contact frequency maps of the W194A-230 system. (A) Representative transmembrane conformations of Cluster 1 to Cluster 4, obtained through distance-based clustering of molecular dynamics trajectories. The structural models display varied spatial arrangements of the TREM2-DAP12 complex in the absence of the W194 residue, illustrating shifts in helix packing and orientation. (B) Corresponding contact frequency maps between TREM2 and DAP12 Chain A and Chain B for Clusters 1-4. Frequency values range from low (black) to high (white), with intermediate contacts shown in shades of red.

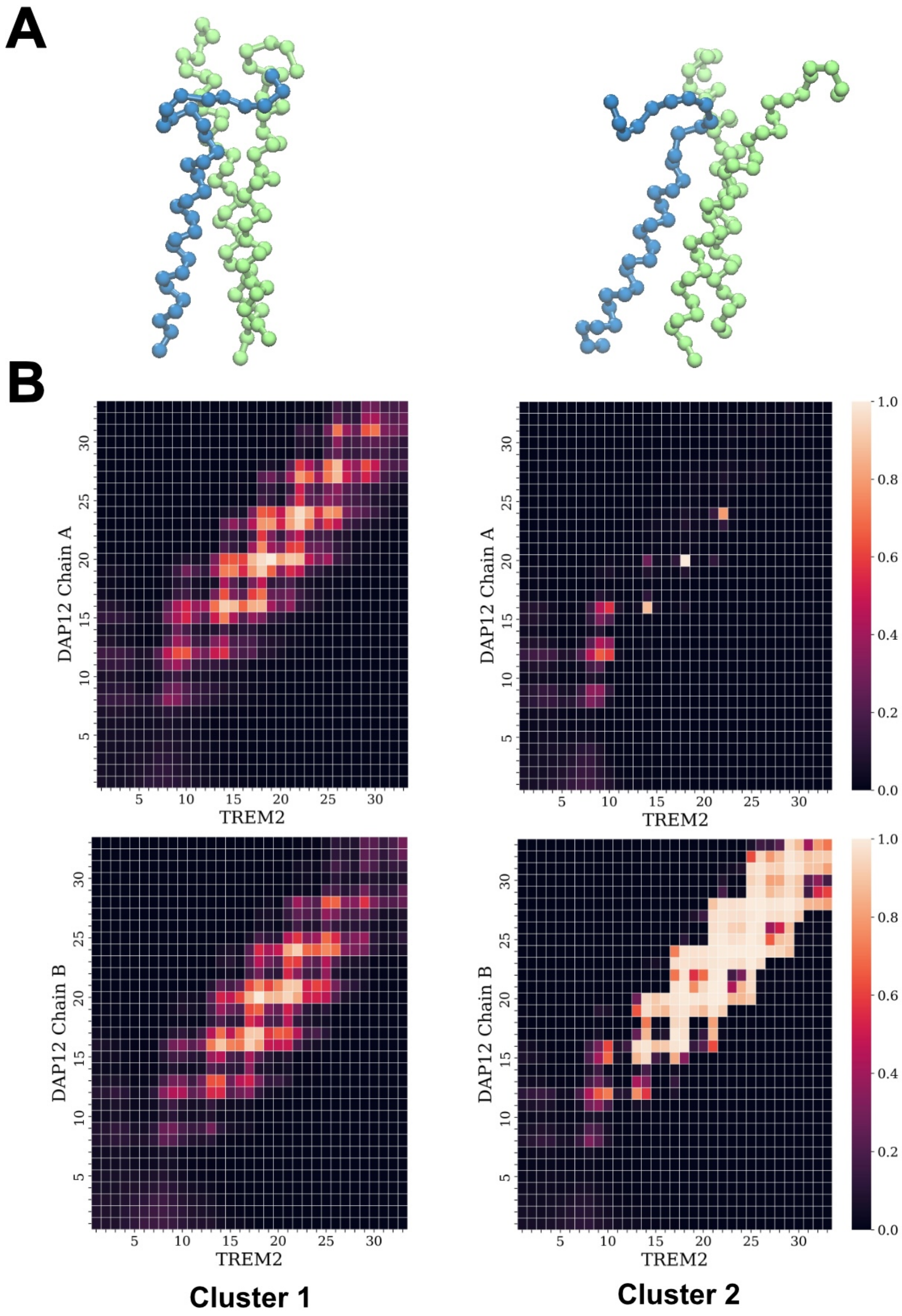


Figure S5: Structural ensembles and contact frequency maps of the W194X-230 system. (A) Representative transmembrane conformations of Cluster 1 and Cluster 2, identified through distance-based clustering of MD trajectories. The snapshots highlight altered packing geometries between TREM2 and DAP12 in the absence of the W194 residue, particularly in helix-helix interface positioning. (B) Corresponding contact frequency maps between TREM2 and DAP12 Chain A and Chain B for Clusters 1 and 2. Contact frequencies are colour-coded from low (black) to high (white), revealing asymmetric and cluster-specific interaction patterns. Notably, Cluster 2 shows significantly reduced contacts with Chain A and compensatory enhancement with Chain B.

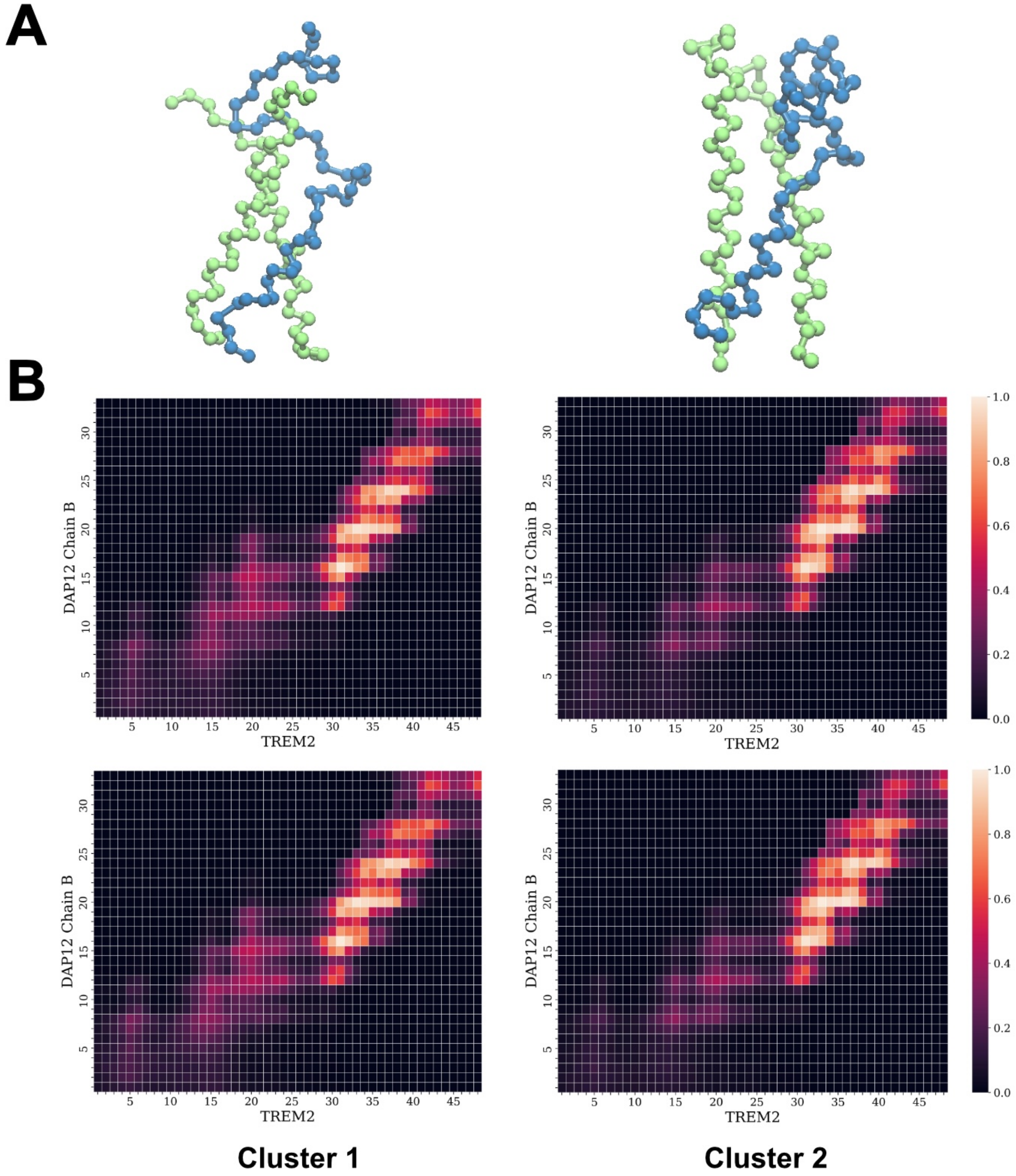


Figure S6: Structural ensembles and contact frequency maps of the WT-219 system. (A) Representative transmembrane conformations of Cluster 1 and Cluster 2, derived from distance-based clustering of MD trajectories. (B) Corresponding contact frequency maps between TREM2 and DAP12 Chain A and Chain B for Clusters 1 and 2. Contact frequencies are color-coded from low (black) to high (white), highlighting consistent and symmetric interaction patterns across both DAP12 chains. Compared to Cluster 1, Cluster 2 exhibits reduced contact frequency in the central transmembrane region.

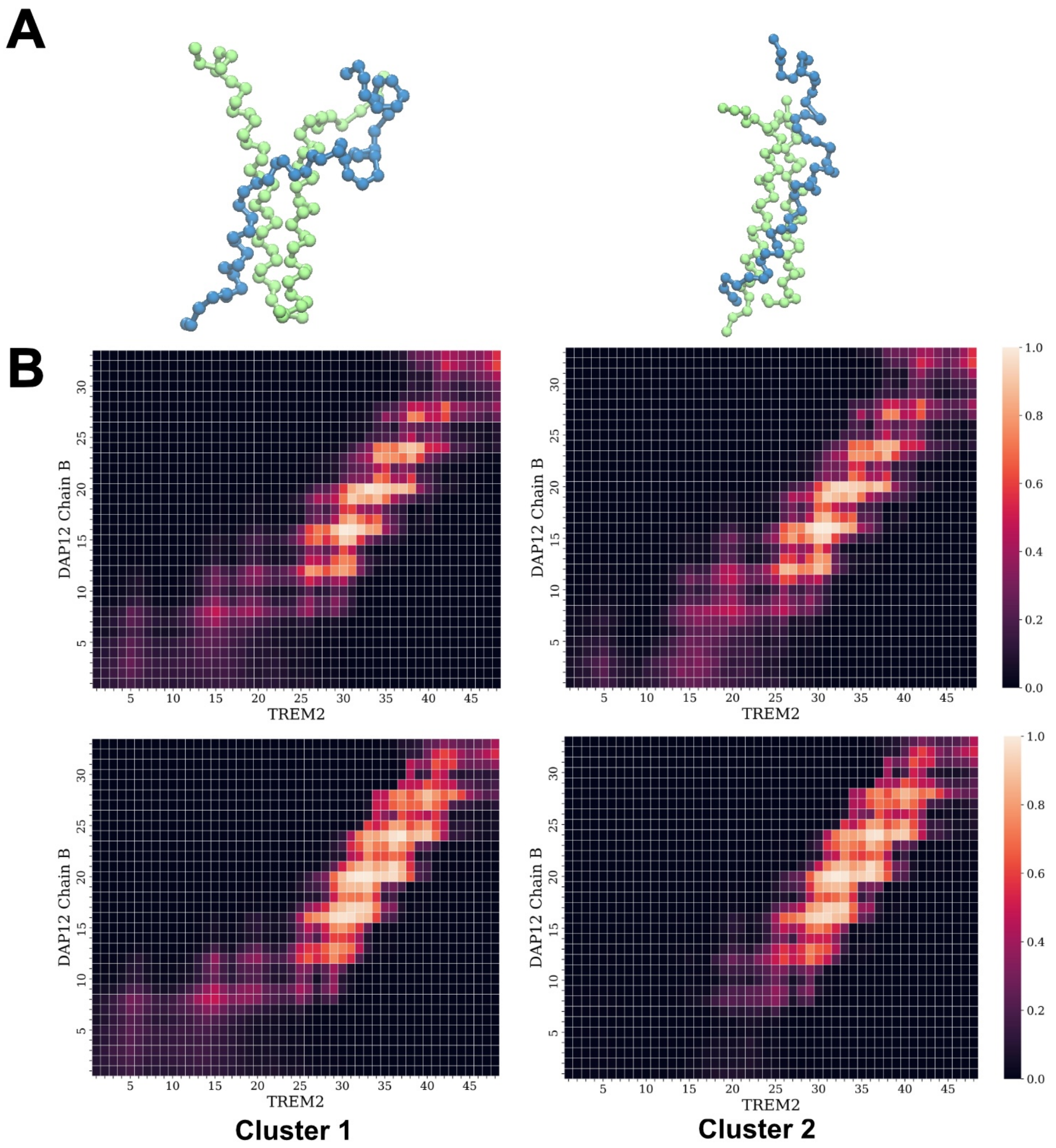


Figure S7: Structural ensembles and contact frequency maps of the W191A-219 system. (A) Representative transmembrane conformations of Cluster 1 and Cluster 2 derived from MD trajectory clustering. (B) Corresponding contact frequency maps between TREM2 and DAP12 Chain A and Chain B for each cluster. Color scale denotes contact frequency from low (black) to high (white).

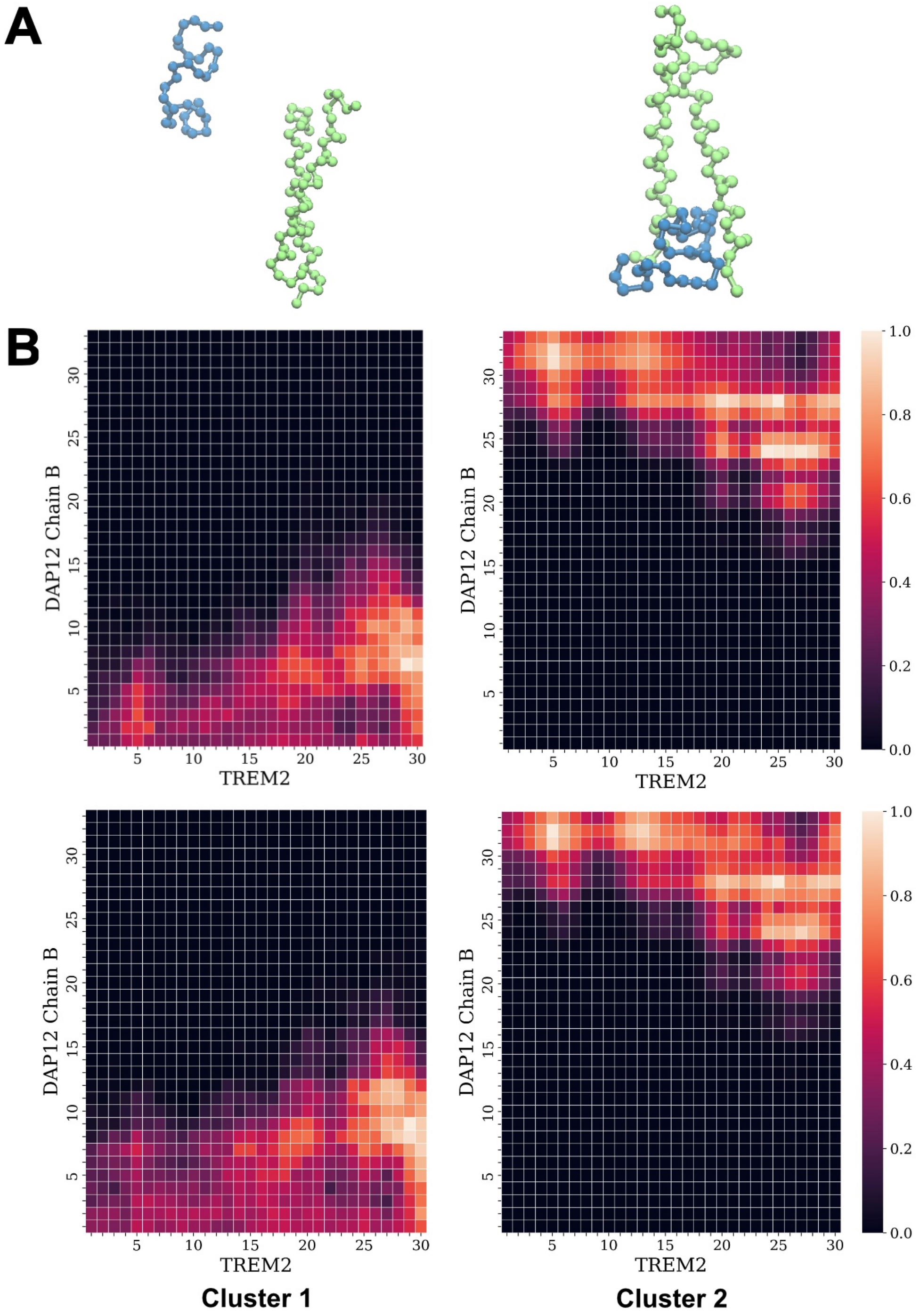


Figure S8: Structural ensembles and contact frequency maps of the W191X-219 system. (A) Representative transmembrane conformations of Cluster 1 and Cluster 2 derived from MD trajectory clustering. (B) Corresponding contact frequency maps between TREM2 and DAP12 Chain A and Chain B for each cluster. Color scale denotes contact frequency from low (black) to high (white). Both Cluster 1 and Cluster 2 display dispersed contact patterns, suggesting a destabilized interface and a shift in binding interactions upon truncation. Notably, Cluster 1 shows contacts between TREM2 and the N-terminal region of DAP12, while Cluster 2 exhibits contacts primarily in the upper transmembrane region.

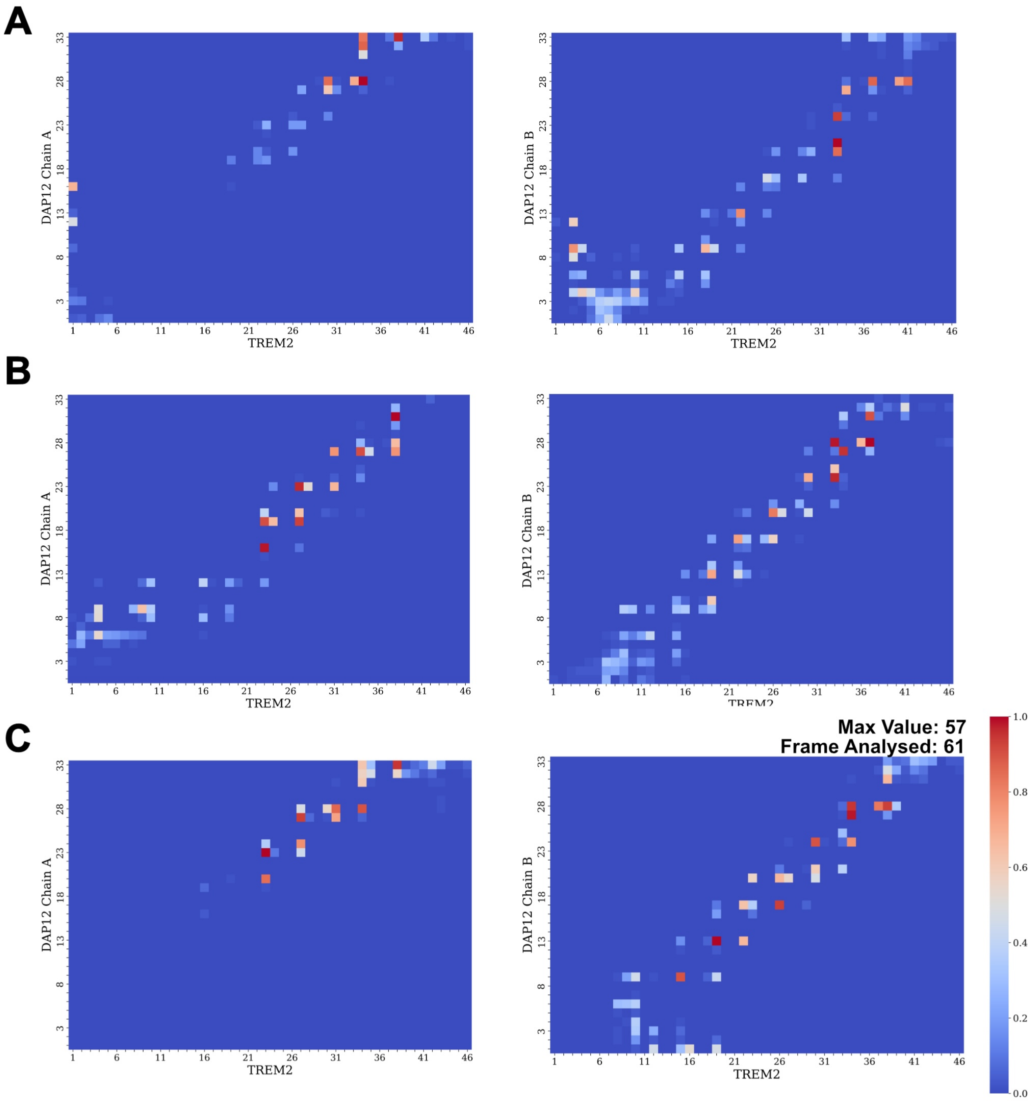


Figure S9: Contact frequency maps of DAP12 with TREM2 in different clusters of the K186A-230 system. (A-C) Contact frequency maps for Cluster 4 (290 $\mu$s), Cluster 1 (295 $\mu$s), and Cluster 3 (298 $\mu$s), respectively. Each panel shows contacts between TREM2 and DAP12 Chain A (left) and Chain B (right). Color scale indicates contact frequency from low (blue) to high (red), normalized to a maximum value of 1.0. The similarity across panels A-C highlights consistent interaction patterns in replicate trajectories of the K186A-230 system.

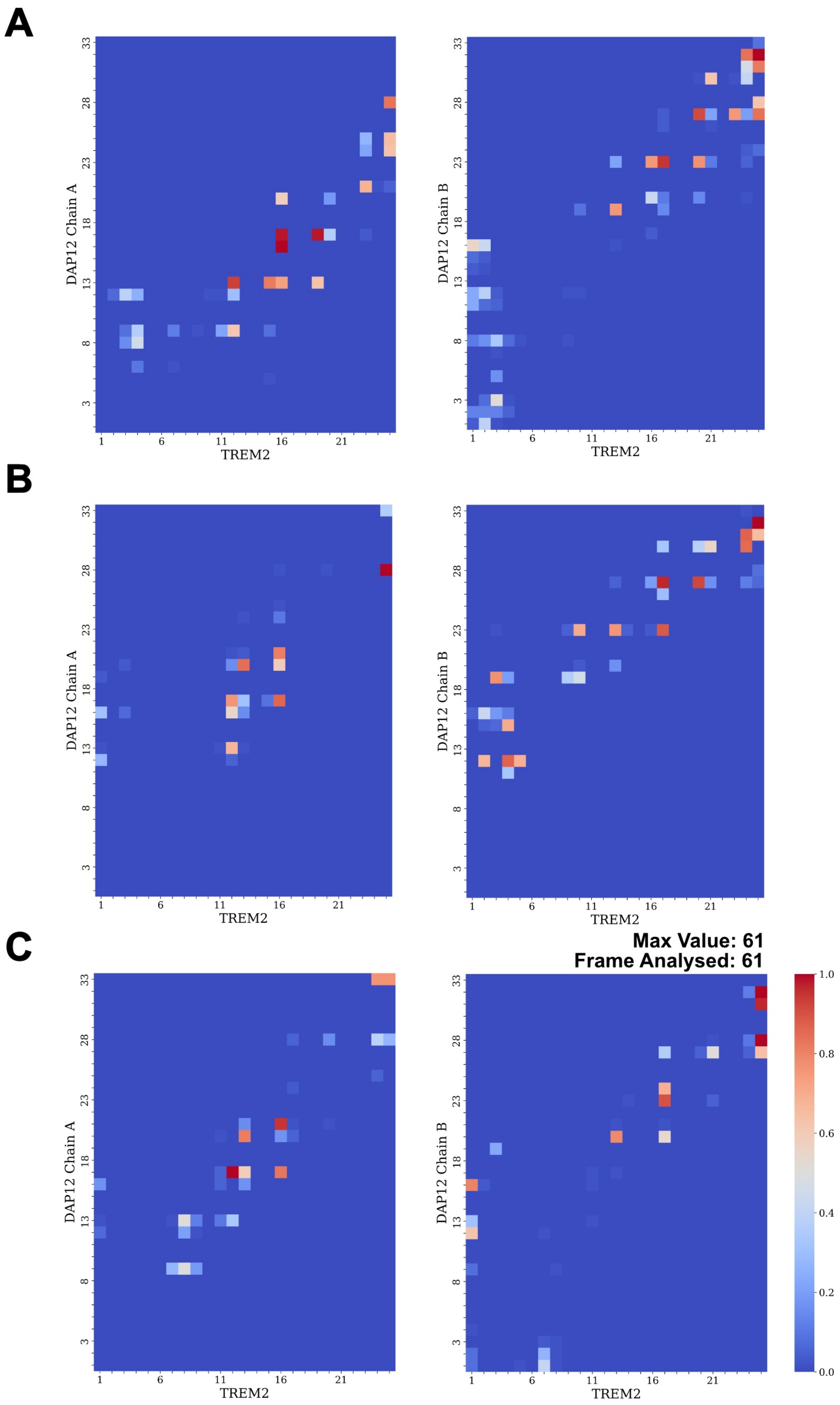


Figure S10: Contact frequency maps of DAP12 with TREM2 in different clusters of the K186X-230 system. (A-C) Contact frequency maps for Cluster 2 (297 $\mu$s), Cluster 4 (299 $\mu$s), and Cluster 4 (300 $\mu$s), respectively. Left and right panels in each subfigure show contact frequencies between TREM2 and DAP12 Chain A and Chain B. The color scale indicates contact frequency, ranging from low (blue) to high (red), normalized to a maximum value of 1.0.

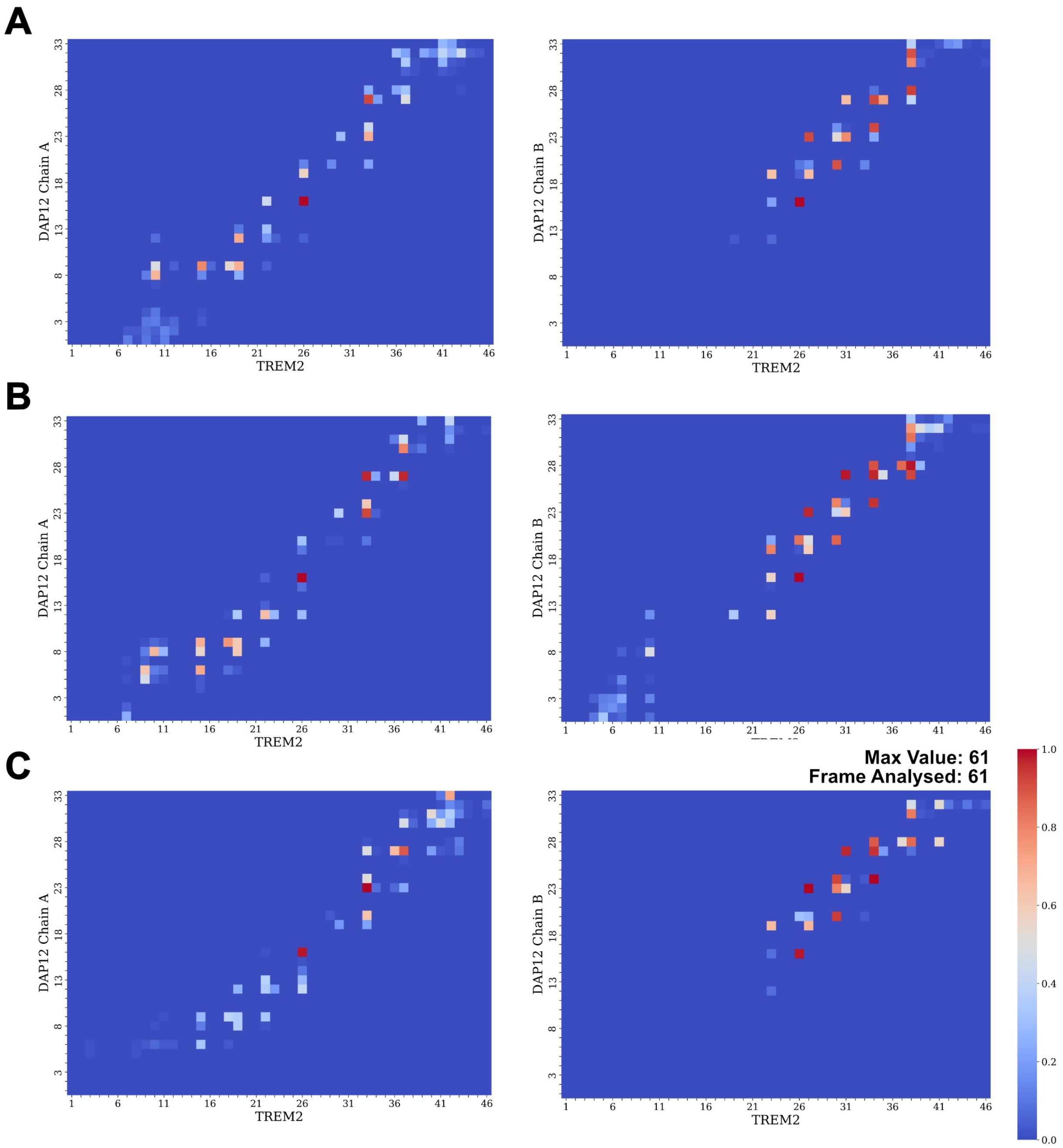


Figure S11: Contact frequency maps of DAP12 with TREM2 in replicate trajectories of the W194A-230 system. (A-C) Contact frequency maps for Cluster 4 at 297 $\mu$s, 299 $\mu$s, and 300 $\mu$s, respectively. Left and right panels in each subfigure show contact frequencies between TREM2 and DAP12 Chain A and Chain B. The color scale represents contact frequency from low (blue) to high (red), normalized to a maximum of 1.0. Consistent patterns across panels A-C indicate reproducible interaction profiles in the W194A-230 system.

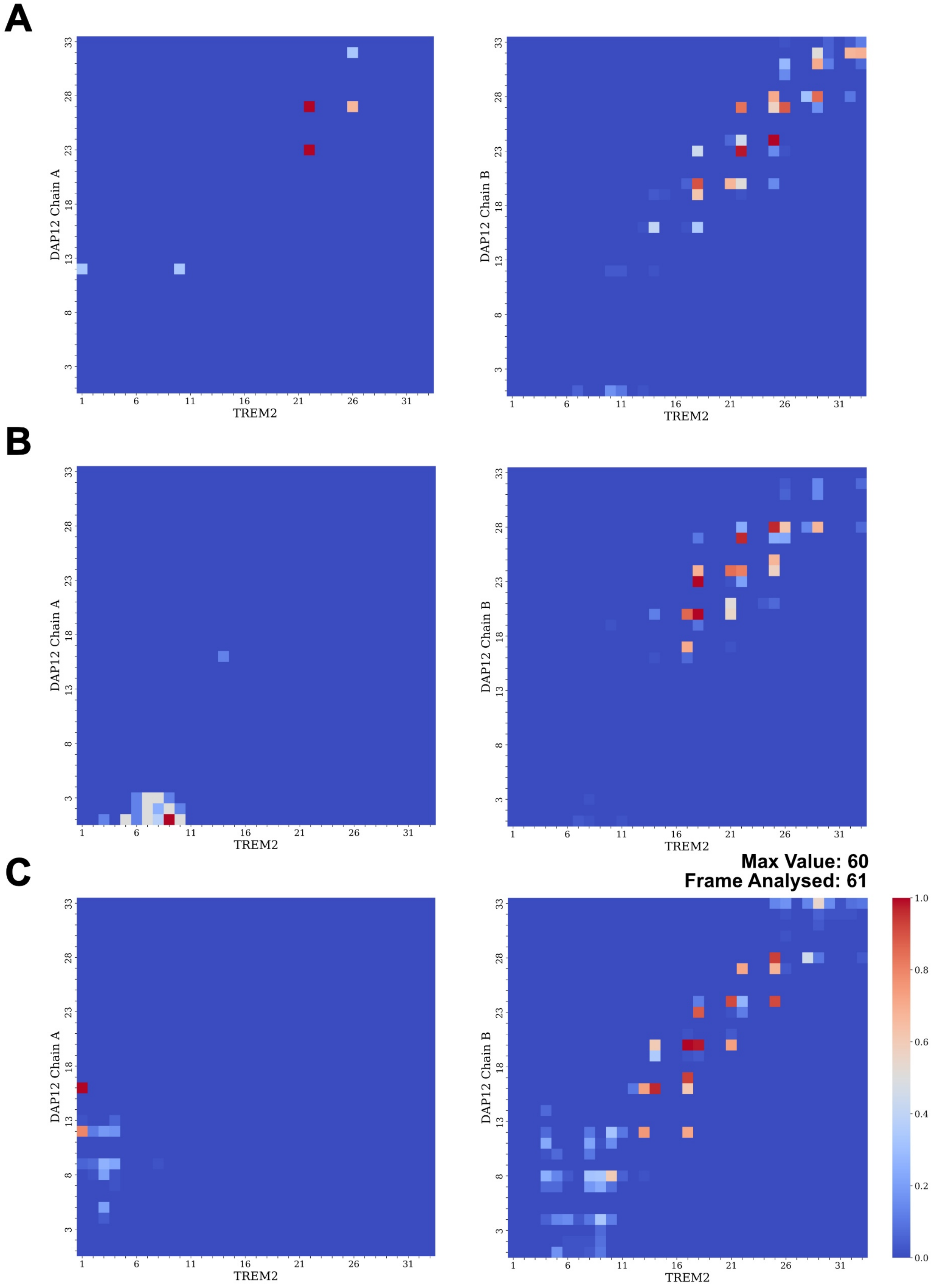


Figure S12: Contact frequency maps of DAP12 with TREM2 in replicate trajectories of the W194X-230 system. (A-C) Contact frequency maps for Cluster 2 at 297 $\mu$s, 299 $\mu$s, and 300 $\mu$s, respectively. Left and right panels in each subfigure show contact frequencies between TREM2 and DAP12 Chain A and Chain B. The color scale represents contact frequency from low (blue) to high (red), normalized to a maximum of 1.0.

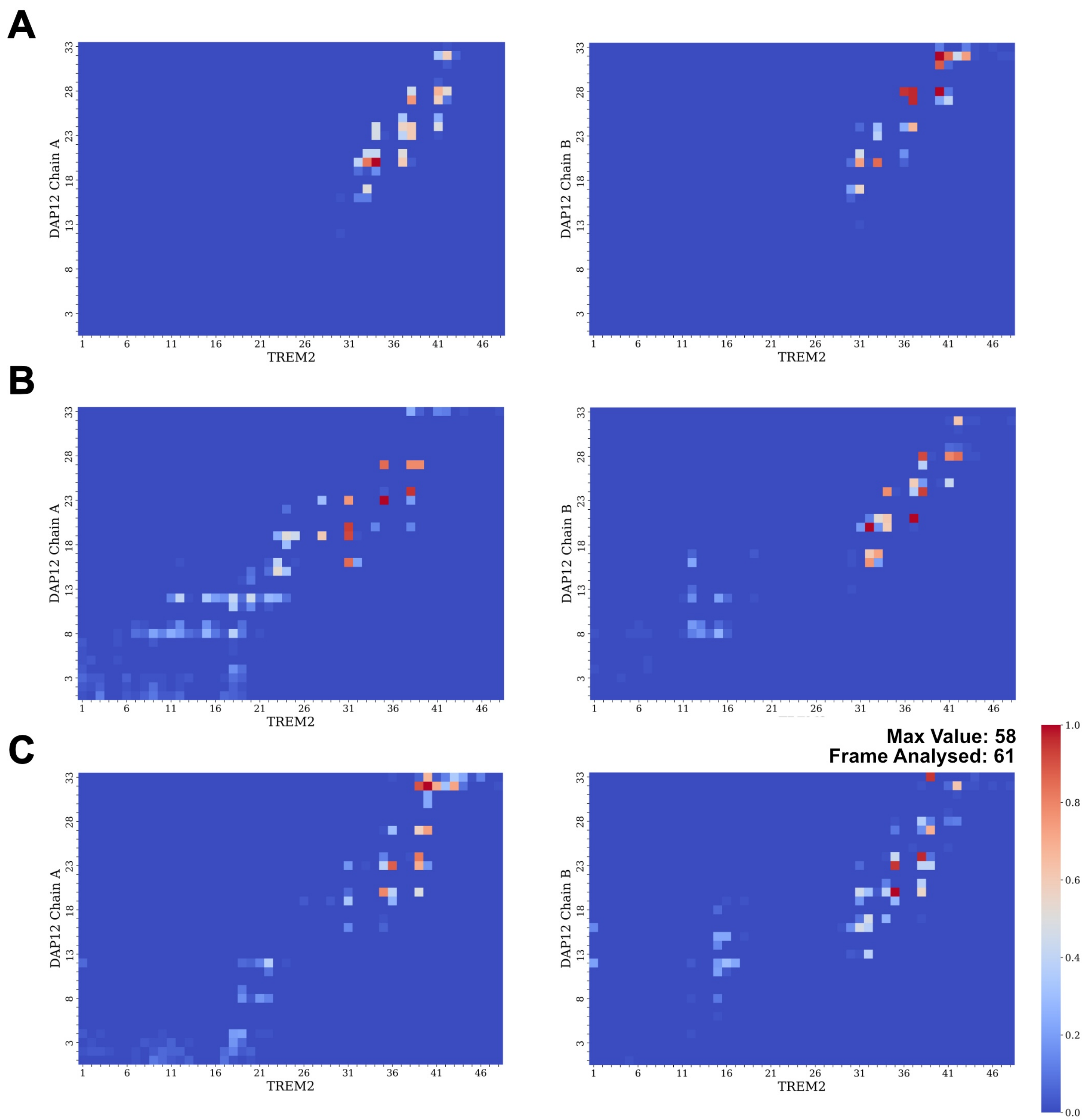


Figure S13: Contact frequency maps of DAP12 with TREM2 in different clusters of the WT-219 system. (A-C) Contact frequency maps for Cluster 1 at 198 $\mu$s, and Cluster 2 at 199 $\mu$s and 200 $\mu$s, respectively. Left and right panels in each subfigure represent contact frequencies between TREM2 and DAP12 Chain A and Chain B. Color scale represents contact frequency from low (blue) to high (red), normalized to a maximum of 1.0.

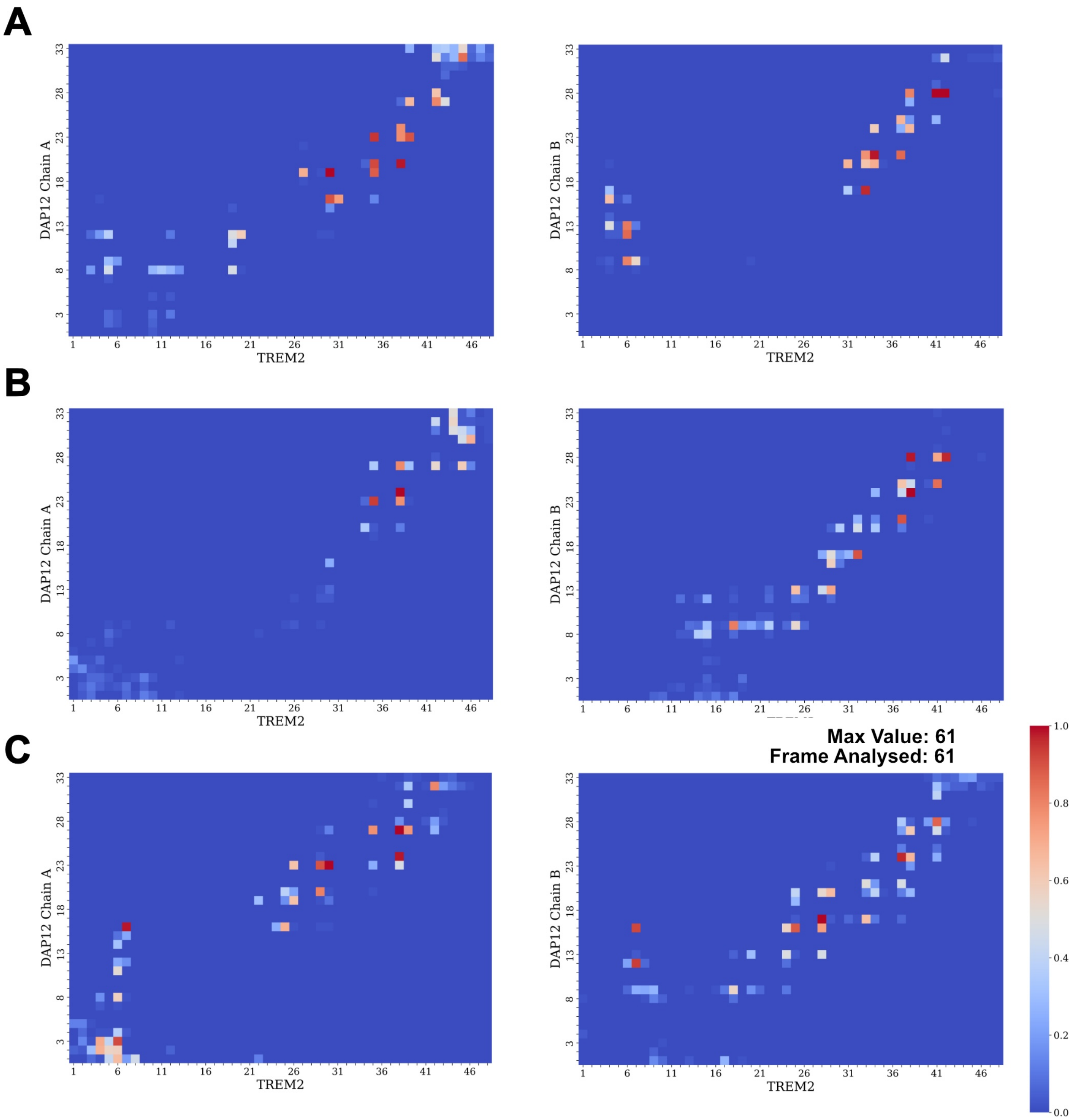


Figure S14: Contact frequency maps of DAP12 with TREM2 in different clusters of the W191A-219 system. (A-C) Contact frequency maps for Cluster 2 at 198 $\mu$s, and Cluster 1 at 199 $\mu$s and 200 $\mu$s, respectively. Left and right panels in each subfigure show contact frequencies between TREM2 and DAP12 Chain A and Chain B. The color scale represents contact frequency from low (blue) to high (red), normalized to a maximum of 1.0.

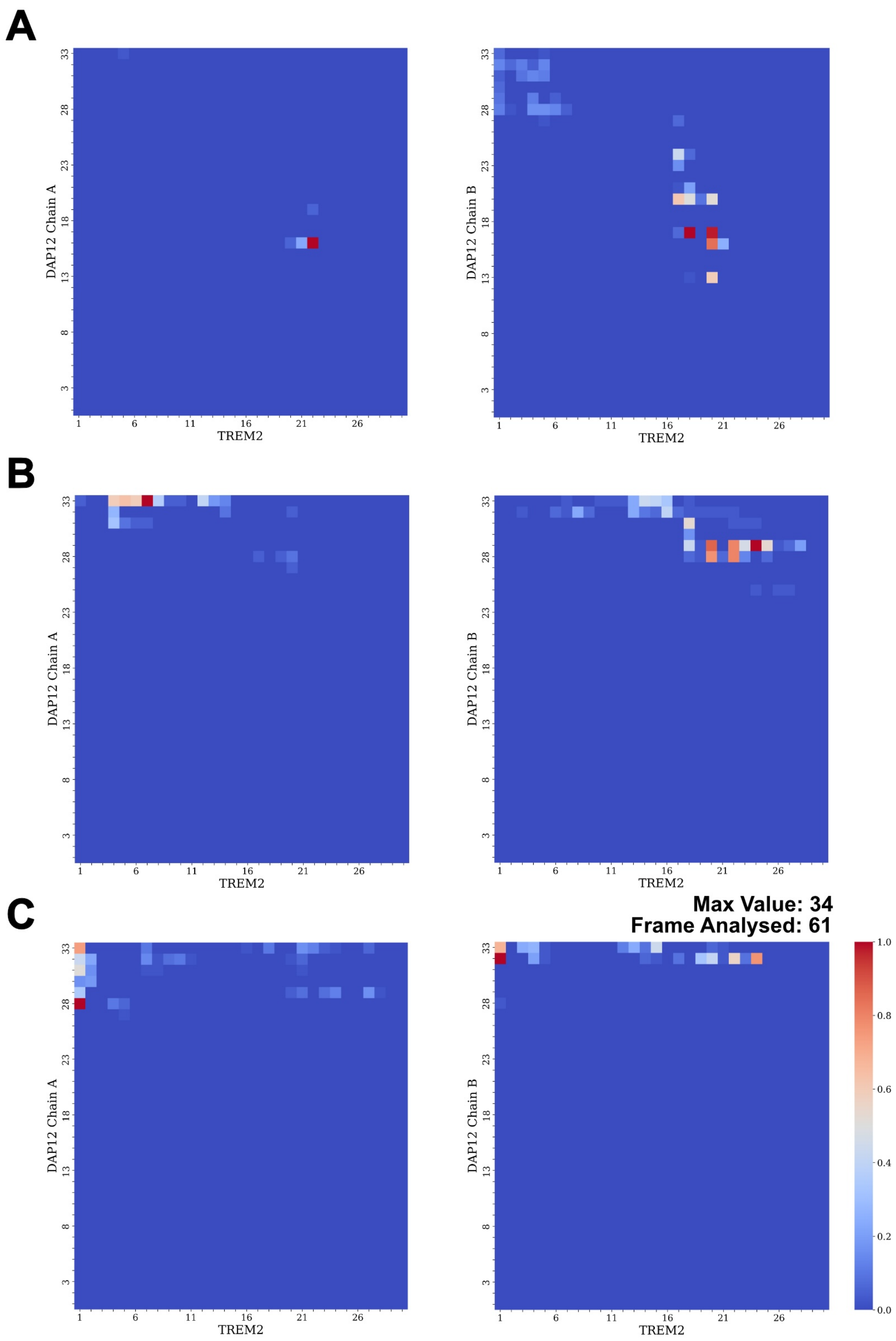


Figure S15: Contact frequency maps of DAP12 with TREM2 in different replicates of the W191X-219 system, all corresponding to Cluster 2. (A-C) Contact frequency maps for simulations at 197 $\mu$s, 198 $\mu$s, and 200 $\mu$s, respectively. Left and right panels in each subfigure display contact frequencies between TREM2 and DAP12 Chain A and Chain B. The color scale represents contact frequency from low (blue) to high (red), normalized to a maximum of 1.0.

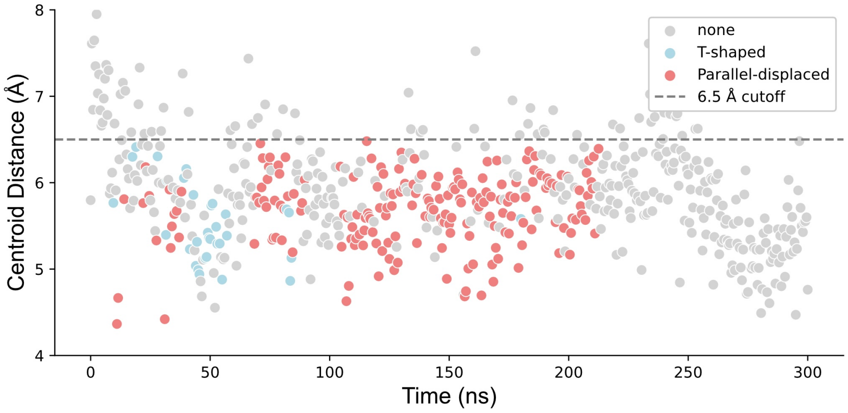


Figure S16: **Hydrogen bonding between TREM2 K186 and DAP12 D50 is maintained in the W194A-230 mutant.** The number of hydrogen bonds ($N_{\mathrm{HB}}$) formed between the side chain of K186 (TREM2) and D50 (DAP12) over time in the W194A-230 mutant. A stable hydrogen bond network is observed throughout the 300 ns simulation: two hydrogen bonds occur in 53.2% of frames, and one bond in 39.1% of frames, with bonding present in 92.3% of the total trajectory. These data support the persistence of this key interfacial interaction in stabilizing the TREM2-DAP12 complex in the W194A-230 mutant.

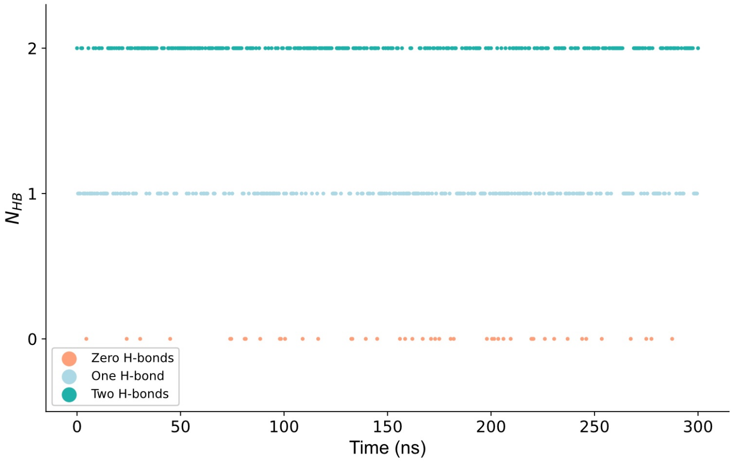


Figure S17: **Time-resolved quantification of hydrogen bonds between K186 of TREM2 and D50 of DAP12 in the W194A-230 mutant system.** Each point represents the number of hydrogen bonds ($N_{\mathrm{HB}}$) formed at a specific time frame across the 300 ns simulation. Hydrogen bonding is categorized into three states: zero (orange), one (light blue), and two (teal) hydrogen bonds. This persistent bonding supports the formation of a tightly packed trimer interface.

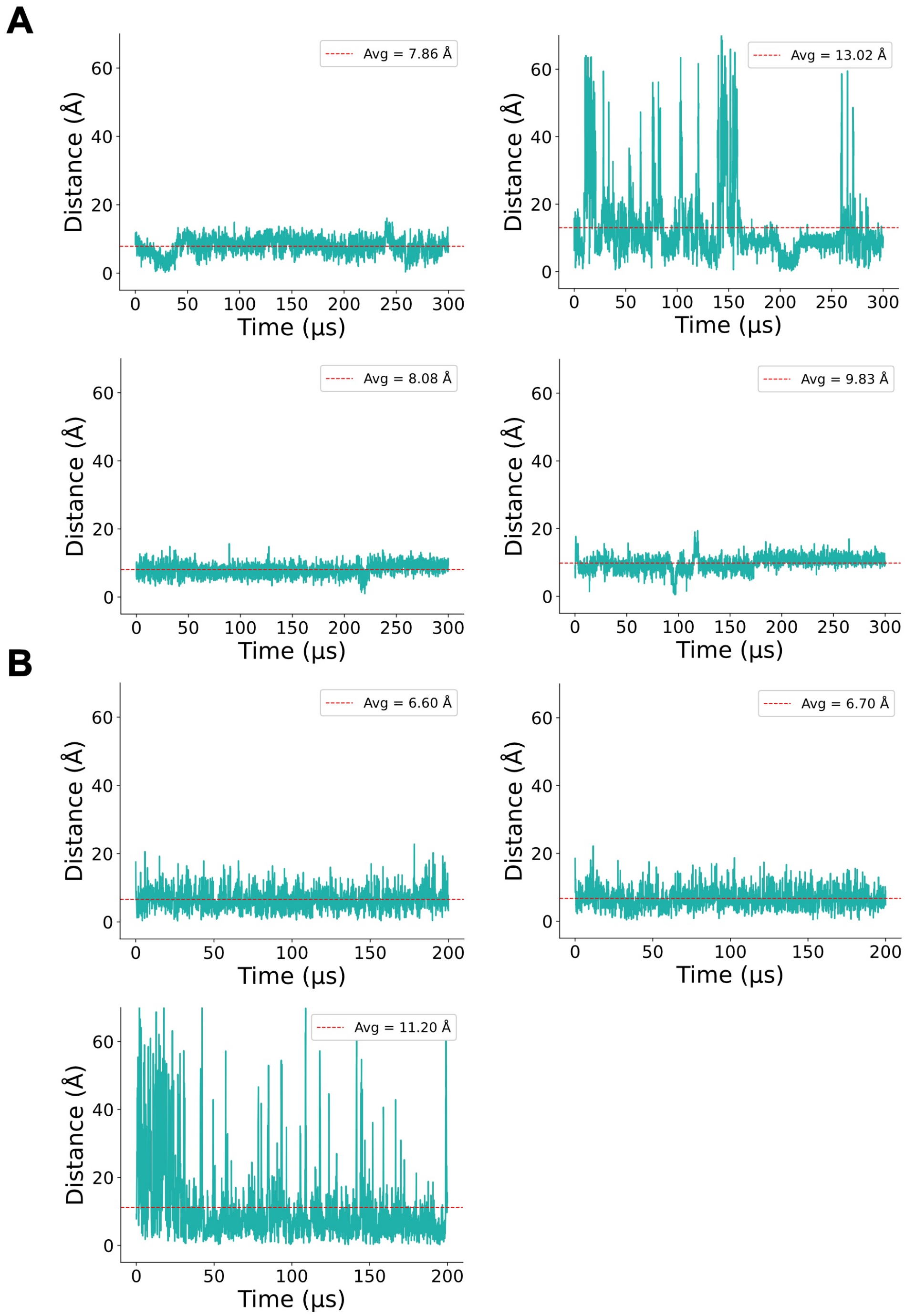


Figure S18: **Centre-of-mass (COM) distance analysis reveals destabilised interfaces in specific TREM2-DAP12 mutants. (A)** COM distances between TREM2 and DAP12 in the 230-residue isoform for K186A, K186X, W191A, and W191X mutants. The K186X-230 mutant shows a markedly higher average distance (13.02 Å), suggesting a disrupted or weakened interface, whereas K186A-230, W194A-230, and W194X-230 maintain shorter average distances (7.86 Å, 8.08 Å, and 9.83 Å, respectively). **(B)** COM distances for the 219-residue isoform in WT, W191A, and W191X constructs. WT-219 and W191A-219 maintain tight packing (6.60 Å and 6.70 Å), while W191X-219 shows significantly increased spatial separation (11.20 Å).

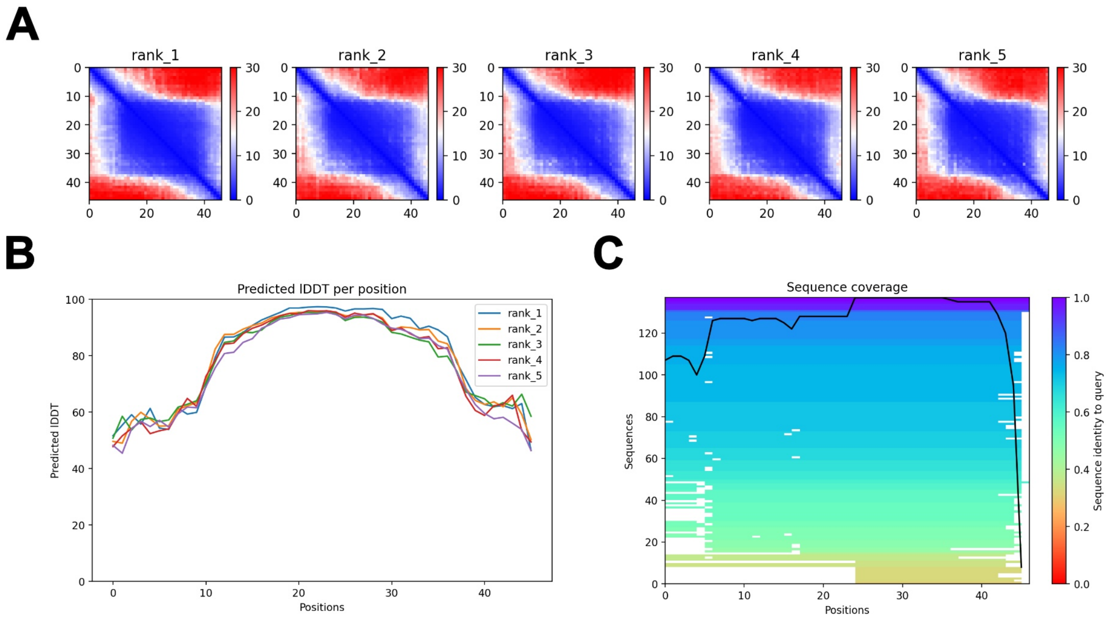


Figure S19: **ColabFold prediction confidence for the TREM2 K186A-230 mutant.** **(A)** Predicted aligned error (PAE) plots for five AlphaFold2 models (rank_1 to rank_5). **(B)** Predicted IDDT scores per residue for each model. **(C)** Multiple sequence alignment (MSA) coverage and identity plotted across the K186A sequence. The black trace indicates overall sequence coverage.

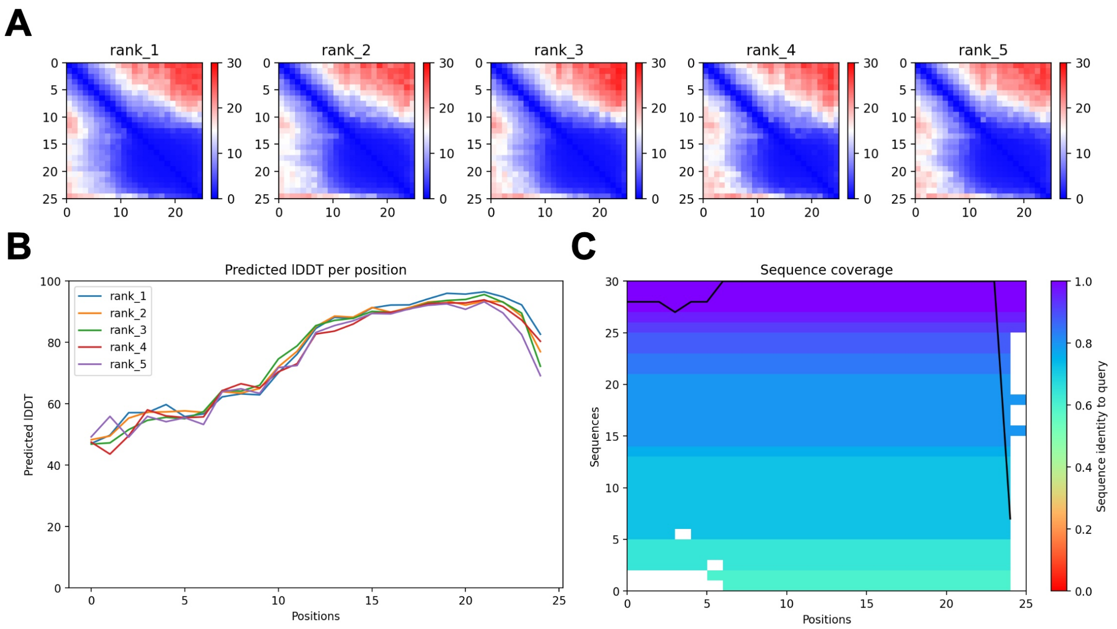


Figure S20: **ColabFold prediction confidence for the TREM2 K186X-230 mutant.** **(A)** Predicted aligned error (PAE) plots for five AlphaFold2 models (rank_1 to rank_5). **(B)** Predicted IDDT scores per residue for each model. **(C)** Multiple sequence alignment (MSA) coverage and identity plotted across the K186X sequence. The black trace indicates overall sequence coverage.

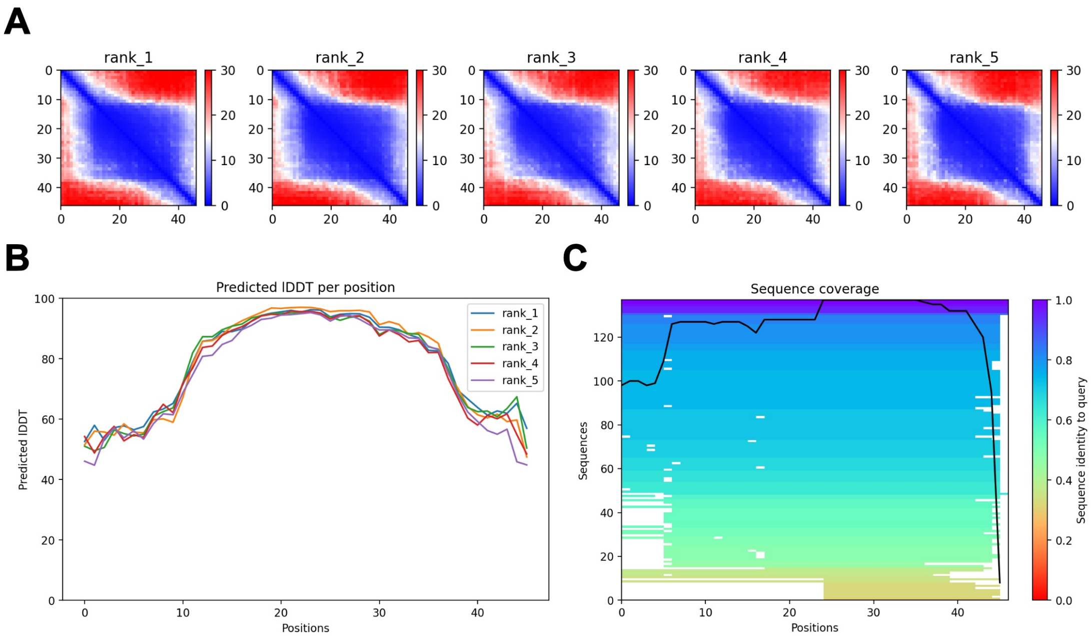


Figure S21: **ColabFold prediction confidence for the TREM2 W194A-230 mutant. (A)** Predicted aligned error (PAE) plots for five AlphaFold2 models (rank_1 to rank_5). **(B)** Predicted IDDT scores per residue for each model. **(C)** Multiple sequence alignment (MSA) coverage and identity plotted across the K186X sequence. The black trace indicates overall sequence coverage.

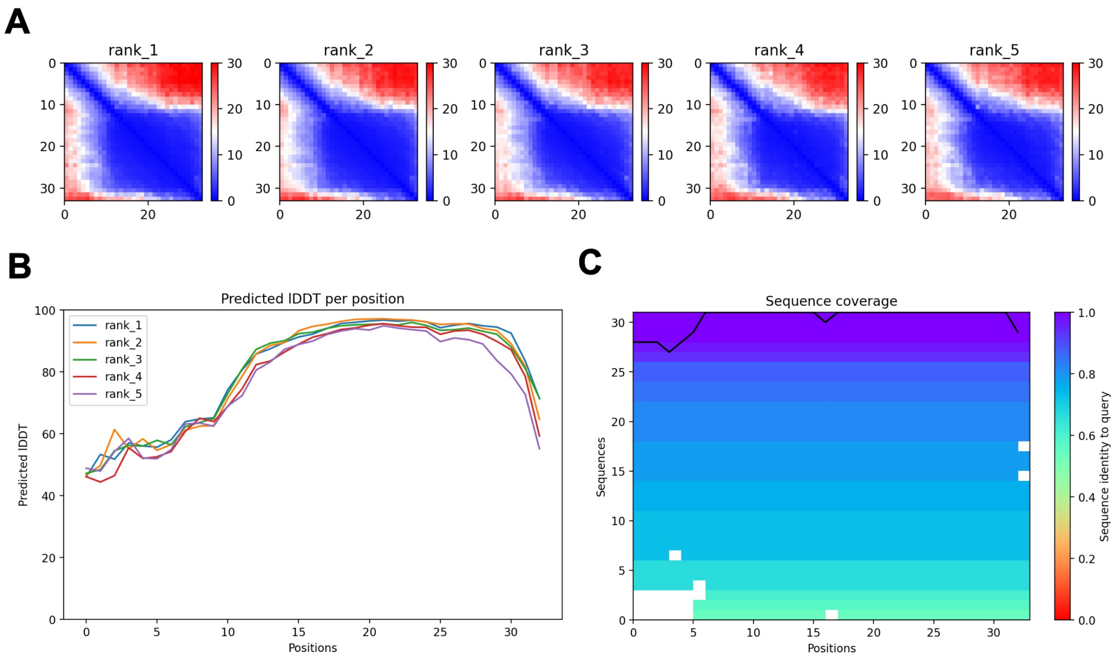


Figure S22: **ColabFold prediction confidence for the TREM2 W194X-230 mutant.** **(A)** Predicted aligned error (PAE) plots for five AlphaFold2 models (rank_1 to rank_5). **(B)** Predicted IDDT scores per residue for each model. **(C)** Multiple sequence alignment (MSA) coverage and identity plotted across the K186X sequence. The black trace indicates overall sequence coverage.

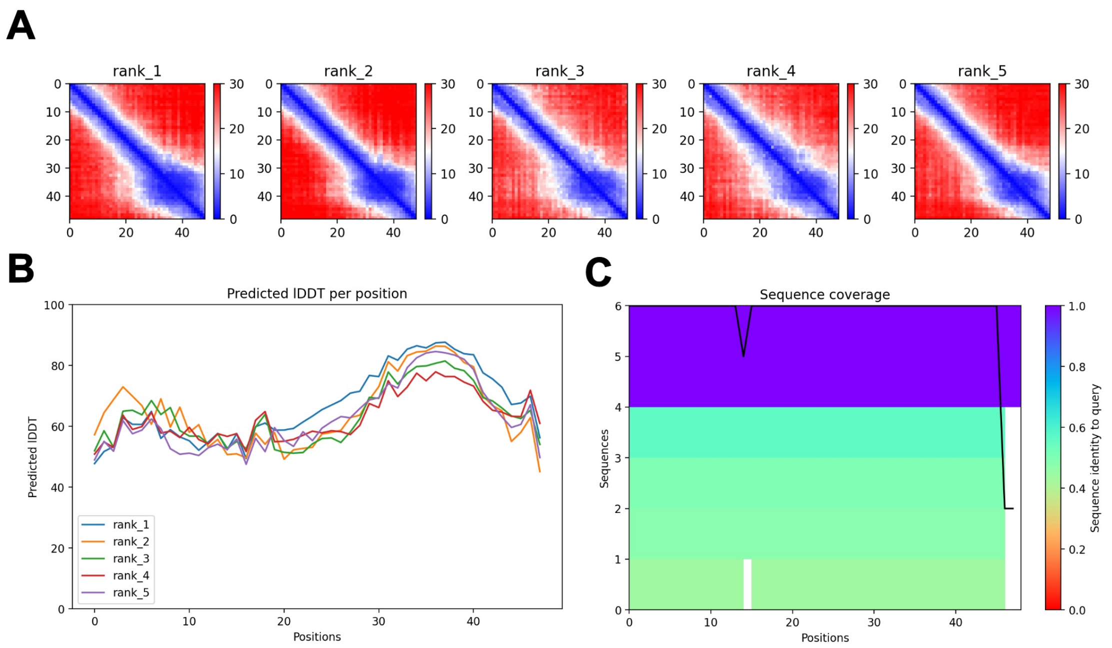


Figure S23: **ColabFold prediction confidence for the TREM2 WT-219 mutant. (A)** Predicted aligned error (PAE) plots for five AlphaFold2 models (rank_1 to rank_5). **(B)** Predicted lDDT scores per residue for each model. **(C)** Multiple sequence alignment (MSA) coverage and identity plotted across the K186X sequence. The black trace indicates overall sequence coverage.

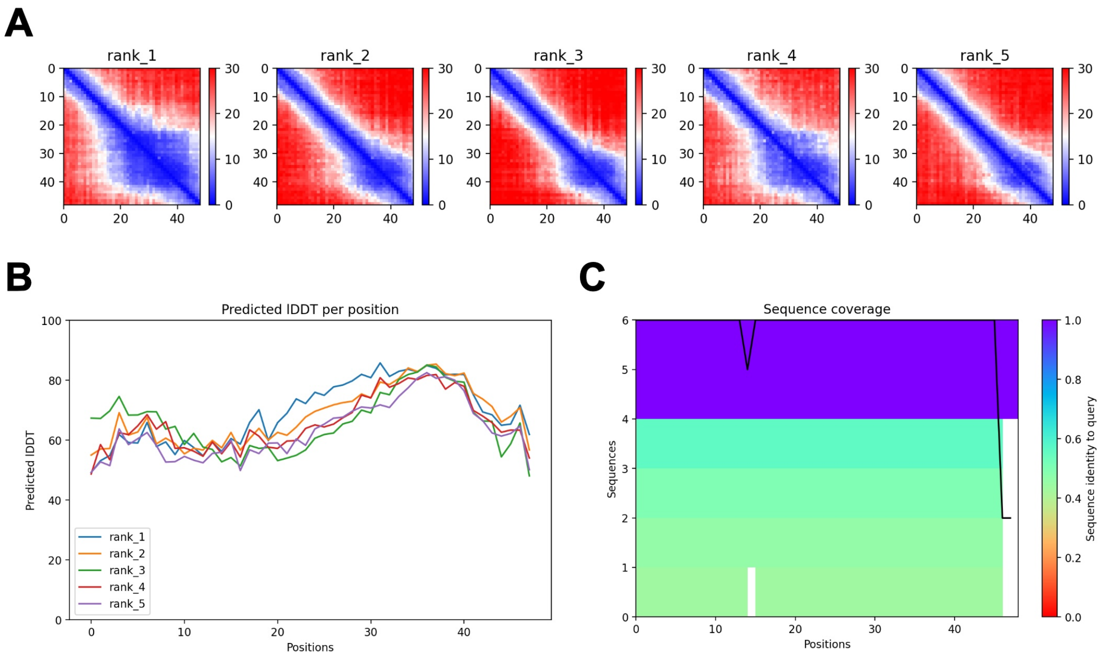


Figure S24: **ColabFold prediction confidence for the TREM2 W191A-219 mutant.** **(A)** Predicted aligned error (PAE) plots for five AlphaFold2 models (rank_1 to rank_5). **(B)** Predicted IDDT scores per residue for each model. **(C)** Multiple sequence alignment (MSA) coverage and identity plotted across the K186X sequence. The black trace indicates overall sequence coverage.

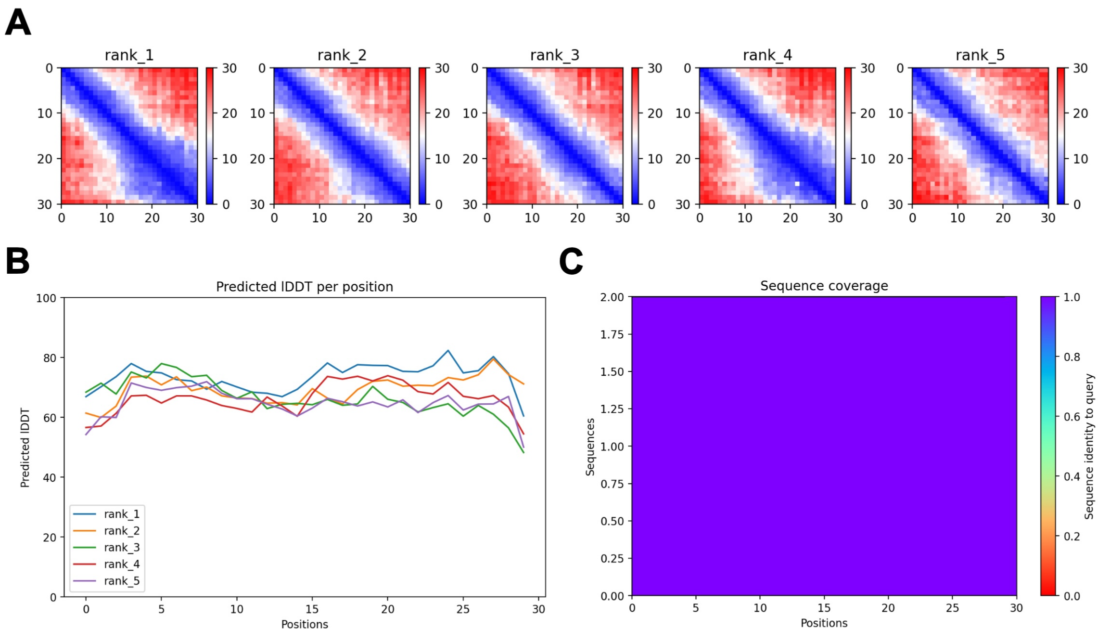


Figure S25: **ColabFold prediction confidence for the TREM2 W191X-219 mutant.** **(A)** Predicted aligned error (PAE) plots for five AlphaFold2 models (rank_1 to rank_5). **(B)** Predicted lDDT scores per residue for each model. **(C)** Multiple sequence alignment (MSA) coverage and identity plotted across the K186X sequence. The black trace indicates overall sequence coverage.

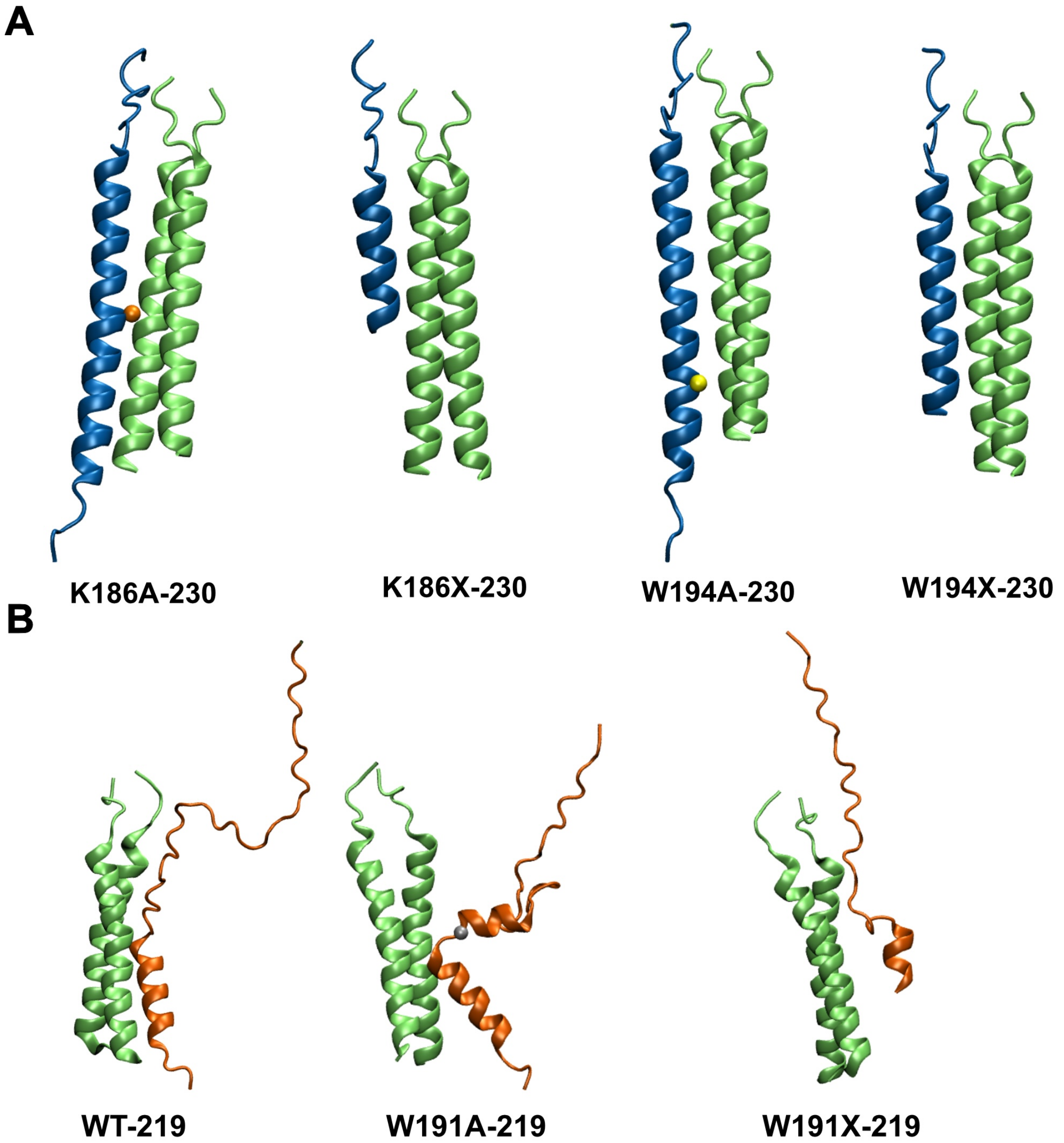


Figure S26: **Initial structures of the TREM2-DAP12 complex for each simulation system. (A)** ColabFold-predicted models of TREM2 transmembrane helix variants (residues 186-230 or 194-230) in complex with DAP12. Mutations include K186A, K186X, W194A, and W194X. TREM2 is shown in green, DAP12 in blue, and the mutation sites are highlighted with spheres. **(B)** Initial configurations of wild-type (WT-219), W191A-219, and W191X-219 systems. TREM2 is shown in green, DAP12 in orange, and mutation sites are highlighted.

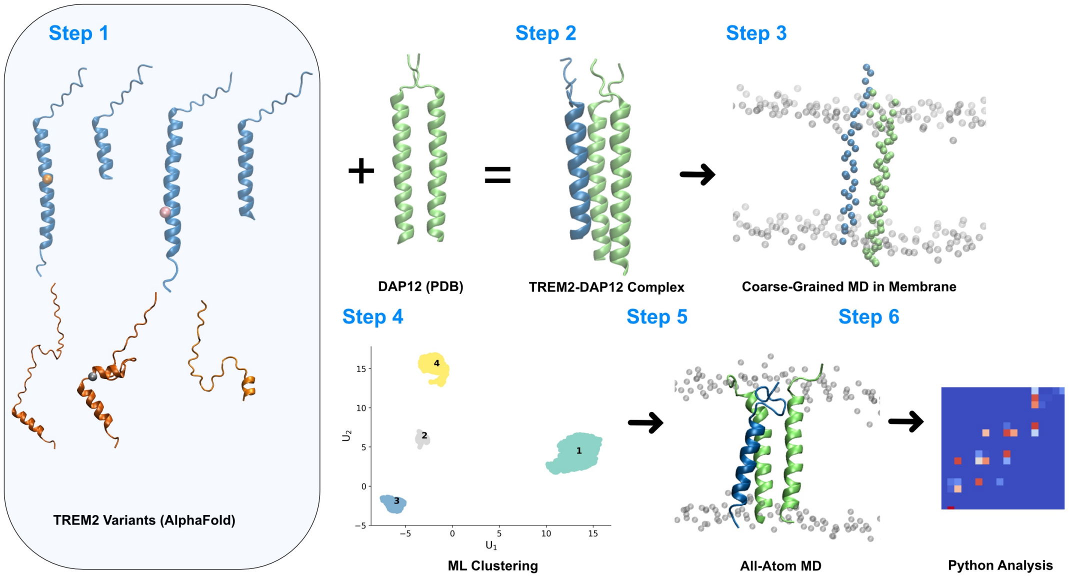


Figure S27: **Computational workflow for investigating TREM2-DAP12 transmembrane complex stability. Step 1:** Structural models of TREM2 isoform 1 and isoform 2 variants were predicted using AlphaFold. **Step 2:** Each TREM2 variant was assembled with DAP12 (PDB: 2L34) to form the initial complex. **Step 3:** The complexes were embedded in a POPC:cholesterol (80:20) bilayer and simulated using coarse-grained (CG) molecular dynamics. **Step 4:** CG trajectories were analyzed with unsupervised machine learning (ML) clustering to identify representative conformational states. **Step 5:** Selected states were converted to atomistic detail for all-atom MD simulations. **Step 6:** Atomistic trajectories were processed using Python-based structural and statistical analyses to quantify mutation effects.

Table S1: Persistent residue-residue contact frequencies.

| | 60% | 70% | 80% | 90% |
|---|---|---|---|---|
| **K186A-230** | I182/V47, F183/T54, L193/A58, W194/V61, W194/Y62, A197/Y62, W198/L66 | W194/V61, W194/Y62, A197/Y62, W198/R67 | W194/V61, W194/Y62, A197/Y62, | W194/V61, W194/Y62 |
| **K186X-230** | R161/D50, P172/V47, P173/T54, I176/L51, L177/I57, A180/V61, I185/Y62, I185/G65, I185/R66 | P173/T54, I176/L51, L177/I57, I185/Y62, I185/G65, I185/L66 | P173/T54, I176/L51, L177/I57, I185/Y62, I185/L66 | I185/Y62, I185/L66 |
| **W194A-230** | P170/V42, F183/L53, K186/D50, I187/L53, I187/I57, A190/T54, A190/I57, A190/V61, S191/V61, L193/I57, L193/V61, W194/A58, W194/V61, W194/Y62, A197/V61, W198/Y62, W198/G65, W198/R66 | K186/D50, I187/I57, A190/T54, A190/I57, A190/V61, A191/V61, L193/I57, L193/V61, W194/A58, W194/V61, A197/V61, W198/Y62, W198/G65, W198/R66 | K186/D50, I187/I57, A190/T54, A190/I57, A190/V61, L191/V61, L193/I57, W194/A58, W194/V61, W198/Y62, W198/G65 | K186/D50, I187/I57, A190/T54, A194/A58, A194/V61, W198/Y62 |
| **W194X-230** | L177/T54, L178/T54, L178/I57, C181/T54, I182/V61, I185/A58, I185/Y62 | L178/T54, L178/I57, I182/V61, I185/A58, I185/Y62 | L178/T54, I182/V61, I185/A58, I185/Y62 | L178/T54 |
| **FL-219** | W191/T54, G194/T54, L195/I57, L198/A58, L198/V61, L199/V61, W200/R66, Q201/R66, E202/R66 | W191/T54, L195/I57, L198/A58, E202/R66 | L198/A58, E202/R66 | L198/A58 |
| **W191A-219** | K167/D50, L195/I57I, V197/V55, L198/I57, L198/A58, L198/V61, L198/Y62, Q201/Y62, E202/Y62, E202/R66 | V197/V55, L198/I57, L198/A58, L198/V61, L198/Y62, Q201/Y62, E202/Y62, E202/R66 | L198/A58, L198/V61, Q201/Y62, E202/Y62 | L198/A58, E202/Y62 |
| **W191X-219** | None | None | None | None |

| | 20% | 30% | 40% |
|---|---|---|---|
| **W191X-219** | R161/Y62, R161/R66, R161/L67, V178/L51, P180/D50, P180/L51 | R161/Y62, R161/R66, R161/L67 | None |

Table S2: Sequence of three systems.

| **chains** | **sequence** |
|---|---|
| DAP12 Chain A | *CSTVSPGVLAGIVVGDLVLTVLIALAVYFLGRL* |
| DAP12 Chain B | *CSTVSPGVLAGIVVGDLVLTVLIALAVYFLGRL* |
| TREM2-WT-230 | *RSLLEGEIPFPPTSILLLLACIFLIKILAASALWAAAWHGQKPGTH* |
| TREM2-K186A-230 | *RSLLEGEIPFPPTSILLLLACIFLIAILAASALWAAAWHGQKPGTH* |
| TREM2-W194A-230 | *RSLLEGEIPFPPTSILLLLACIFLIKILAASALAAAAWHGQKPGTH* |
| TREM2-W194X-230 | *RSLLEGEIPFPPTSILLLLACIFLIKILAASAL* |
| TREM2-WT-219 | *AERHVKEDDGRKSPGEVPPGTSPACILATWPPGLLVLLWQETTLPEH* |
| TREM2-W191A-219 | *AERHVKEDDGRKSPGEVPPGTSPACILATAPPGLLVLLWQETTLPEH* |
| TREM2-W191X-219 | *AERHVKEDDGRKSPGEVPPGTSPACILAT* |
